# EndoMT: An R Package Resource for Endothelial-Mesenchymal Transition Gene Set Enrichment Analysis

**DOI:** 10.64898/2026.09.29.755303

**Authors:** Katie Best, John Casement, Penny E Lovat, Laura Walker, Jane Armstrong

**Affiliations:** Translational and Clinical Research Institute, Newcastle University, Newcastle upon Tyne, United Kingdom; Bioinformatics Support Unit, Newcastle University, Newcastle upon Tyne, United Kingdom; John Dawson Drug Discovery and Development Research Institute, University of Sunderland, Sunderland, United Kingdom

**Keywords:** Endothelial to mesenchymal transition_1_, EndoMT_2_, Gene Set Enrichment Analysis (GSEA)_3_, R package_4_, Transcriptomics_5_

## Abstract

**Introduction:** Endothelial-mesenchymal transition (EndoMT) is a dynamic endothelial cell-state transition increasingly implicated in fibrosis, vascular dysfunction, inflammation and cancer. Despite expansion of EndoMT-focused transcriptomic research, EndoMT representation remains limited to broader epithelial-mesenchymal transition (EMT) frameworks in standard gene set repositories.

**Methods:** To address the absence of a general-purpose EndoMT gene set for transcriptomic enrichment analysis, we developed an open EndoMT gene set resource and accompanying R package. A literature-informed EndoMT gene set was curated and directionally assigned to EndoMT UP and EndoMT DOWN signatures according to reported or hypothesised expression during partial EndoMT, defined here as an active transitional state rather than a complete mesenchymal endpoint, to improve capture of EndoMT-associated transcriptional changes across different stages of the process. Genes were annotated by marker subtype and implemented as Gene Matrix Transposed files distributed through the R package, together with exported R objects and gene-level metadata. Benchmarking was performed using bulk RNA-seq data sets GSE118446, comprising seven-day IL-1β/TGF-β2-treated human umbilical vein endothelial cells (HUVECs) and human pulmonary arterial endothelial cells, and GSE342578, comprising three-day TGF-β1/2/3-treated HUVECs; single-cell RNA-seq data set GSE237895, comprising seven-day TGF-β1-treated and SNAI1-overexpressing HUVECs; and endothelial cell and cancer-associated fibroblast populations from human metastatic melanoma single-cell RNA-seq data set GSE72056.

**Results:** In GSE118446, EndoMT UP was positively enriched and EndoMT DOWN negatively enriched in chronically IL-1β/TGF-β2-treated endothelial cells, concordant with experimentally validated EndoMT induction, while partial EndoMT-associated enrichment was detected in single ligand-treatment. In contrast, EndoMT UP was negatively enriched following acute TGF-β1/2/3 treatment, suggesting temporal differences between acute TGF-β stimulation and chronic EndoMT induction. In GSE237895 mesenchymal-like endothelial cells, EndoMT UP was positively enriched and EndoMT DOWN negatively enriched in both TGF-β1-treated and SNAI1-overexpressing models. EndoMT gene sets also discriminated endothelial and mesenchymal/stromal transcriptional states in GSE72056 metastatic melanoma. Marker subtype and Gene Ontology Biological Process analyses provided additional exploratory resolution of EndoMT-associated components beyond MSigDB Hallmark EMT.

**Discussion:** The EndoMT R package provides a dedicated, accessible and interoperable resource for EndoMT-focused transcriptomic enrichment analysis. Benchmarking demonstrates utility for detecting established, partial and divergent EndoMT-associated states, supporting broad applicability across EndoMT biology research.

## 1 Introduction

Endothelial-mesenchymal transition (EndoMT) is a dynamic cell-state transition characterised by loss of endothelial markers and acquisition of mesenchymal characteristics. EndoMT is a spectrum and may be classified as complete, with complete loss of endothelial lineage gene expression and high levels of expression of mesenchymal-specific genes, or partial, a potentially transient and reversible state of plasticity in which endothelial lineage markers are reduced but not fully lost (Simons, 2023; Yang *et al*., 2020). EndoMT was first reported in embryonic development as an essential mechanism for normal cardiac morphogenesis, specifically the cardiac cushion mesenchyme which subsequently remodels to form the cardiac valves and septum (Barnett and Desgrosellier, 2003; Liu and Nakano, 2024; Markwald, Fitzharris and Manasek, 1977; Markwald, Fitzharris and Smith, 1975; Person, Klewer and Runyan, 2005). Similar to the more extensively studied process of epithelial-mesenchymal transition (EMT), EndoMT involves mesenchymal transdifferentiation but is specific to endothelial cells (Medici *et al*., 2010; Qian *et al*., 2025). Endothelial cells are a functionally and phenotypically heterogenous mesoderm-derived cell population with arterial, venous or lymphatic fates, and properties specific to their destined vascular bed (Cho *et al*., 2018; Dejana, Hirschi and Simons, 2017). They are essential for maintaining vascular homeostasis and barrier function through regulation of tone, permeability and immune recruitment, mediated in part by adhesion molecules and chemokines (Félétou, 2011; Sena, Pereira and Seiça, 2013; Vestweber, 2012; Pober and Sessa, 2007; Sturtzel, 2017). In contrast, mesenchymal cells are stellate or spindle-shaped stromal cells, with transmigratory, contractile and extracellular matrix-producing functions (Qian *et al*., 2025; von Gise and Pu, 2012; Dejana, Hirschi and Simons, 2017).

Accumulating evidence implicates aberrant EndoMT in a broad range of pathological states, including atherosclerosis (Chen *et al*., 2015), neointima formation (Cooley *et al*., 2014), heterotopic ossification (Yung *et al*., 2015), pulmonary hypertension (Arciniegas *et al*., 2007), diabetic nephropathy (Kizu, Medici and Kalluri, 2009; Li, Qu and Bertram, 2009), cerebral cavernous malformations (Maddaluno *et al*., 2013), and cardiac, pulmonary and renal fibrosis (Zeisberg *et al*., 2007b; Zeisberg *et al*., 2008). In cancer, EndoMT has been linked to tumour progression and proposed as a source of cancer-associated fibroblasts (CAFs) (Potenta, Zeisberg and Kalluri, 2008; Zeisberg *et al*., 2007a). CAFs are key mediators of tumour growth and vascular remodelling (Sewell-Loftin *et al*., 2017); they contribute to tumour-supportive environments through extracellular matrix deposition, secretion of tumour-promoting cytokines and pro-angiogenic factors, and release of exosomes in promoting chemoresistance (Saw, Chen and Song, 2022; Bhome *et al*., 2017). While current evidence indicates transforming growth factor-beta (TGF-β) signalling and MEK5/ERK5/BMP pathway activation to be the two most well-characterised drivers of EndoMT (Cuttano *et al*., 2016; Renz *et al*., 2015; Chen *et al*., 2012; Chen *et al*., 2014), alongside Wnt/β-catenin, Notch and inflammatory cascades (Kovacic *et al*., 2012; Liebner *et al*., 2004), a deeper understanding of the molecular signals and regulatory pathways critical to the induction, progression and maintenance of EndoMT remains a subject of active research (Dejana, Hirschi and Simons, 2017). Defining the mechanisms driving this multifaceted process is important for identifying disease-relevant pathways and informing novel EndoMT-targeted therapeutic strategies (Jackson *et al*., 2017; Cho *et al*., 2018). Despite the growing importance of EndoMT in disease and the increased use of transcriptome-scale approaches to study this biological process, there remains a need for a dedicated gene set resource that enable EndoMT-associated transcriptional programmes to be interrogated directly.

In view of the rapid expansion and technological advancement of high-throughput omics, biomedical systems research has increasingly shifted towards the large-scale analysis of transcriptomic, genomic, proteomic and metabolomic data sets (Manzoni *et al*., 2018). Whilst these approaches have been instrumental in enhancing our understanding of health and disease, they also present challenges in how dynamic molecular systems are interrogated, and how biologically relevant functions and interactions are inferred. Gene set- and pathway-based methods therefore provide an important framework for transforming large transcriptomic profiling data sets into biologically meaningful units, including phenotypic signatures, regulatory pathways, cell-cell and ligand-receptor interactions, from which mechanistic insight can be extrapolated (Wanichthanarak, Fahrmann and Grapov, 2015). Gene set repositories, including the Gene Ontology (GO) Consortium (Blake *et al*., 2013), Kyoto Encyclopedia of Genes and Genomes (KEGG) (Kanehisa *et al*., 2025), Reactome (Ragueneau *et al*., 2026) and The Broad Institute’s Human Molecular Signatures Database (MSigDB) (Liberzon *et al*., 2011), are essential bioinformatics resources for the analysis of differential expression results, providing curated collections of predefined gene sets grouped by shared biological function or features, frequently in the Gene Matrix Transposed (GMT) file format (Hagenauer *et al*., 2024). Several algorithms are available for conducting gene-set based analyses in R (Subramanian *et al*., 2005; Wu and Smyth, 2012; Wu *et al*., 2010; Ritchie *et al*., 2015); Gene Set Enrichment Analysis (GSEA) (Subramanian *et al*., 2005) is among the most widely used method in this field, evaluating whether a predefined set of genes show statistically significant, coordinated differences between biological states.

MSigDB (Liberzon *et al*., 2011) is a prominent gene set repository consisting of tens of thousands of annotated gene sets, divided into nine major collections, including the Hallmark (H) collection. The Hallmark collection comprises 50 curated gene sets designed to represent well-defined biological states or processes with coherent expression (Liberzon *et al*., 2015). Historically, EndoMT has been considered a subtype of EMT (Carew, Wang and Kantharidis, 2012). Although EndoMT is now recognised as a distinct biological process (Zeisberg *et al*., 2007b), there is no EndoMT-specific Hallmark gene set or equivalent general-purpose predefined collection. Consequently, EndoMT-associated transcriptional programmes currently remain largely captured under the broader framework of EMT in standard enrichment analyses.

We therefore aimed to curate a bespoke literature-derived EndoMT gene set for human transcriptomic research through review of the literature, encompassing *in vitro* and disease-context studies of EndoMT across endothelial cell types, inflammatory stimuli, metabolic dysfunction and pathological settings. The curated resource was designed to capture endothelial-associated molecular features of EndoMT, including molecular regulators, signalling pathways and phenotypic markers that are not captured within existing EMT-focused gene sets. Here we present directionally assigned EndoMT UP and EndoMT DOWN gene sets provided as GMT files and distributed through an openly accessible R package for transcriptomic GSEA. Directional assignment was centred on partial EndoMT because many EndoMT-relevant genes represent active, context-dependent transitional responses that may occur during progression towards complete EndoMT, or reflect a transient process, and are not necessarily retained at a fully mesenchymal endpoint. This approach was intended to improve applicability across transcriptomic studies spanning different stages of EndoMT activation. We demonstrate the utility of this resource through benchmarking in an established EndoMT model, integration with existing Hallmark GSEA workflows and integrated downstream data visualisation of EndoMT-associated enrichment patterns.

## 2 Methods

### 2.1 Curation of the EndoMT gene sets

A bespoke EndoMT-focused gene set was curated through a comprehensive narrative literature review of more than 300 PubMed-indexed original research and review articles (Supplementary Table 1). Searches were conducted between May 2025 and April 2026 using PubMed (RRID:SCR_004846) and Ovid MEDLINE (RRID:SCR_002185). Search terms included “EndMT” and “endothelial-mesenchymal transition,” as well as broader terms “EMT” and “epithelial-mesenchymal transition”, recognising that “endothelial-mesenchymal transition” was only introduced as a medical subject heading (MeSH, RRID:SCR_004750) descriptor in 2024 (National Library of Medicine, 2026), with all prior EndoMT-related literature indexed under the broader EMT MeSH hierarchy. Searches were refined further using context-specific terms related to angiogenesis, cancer, cardiovascular disease, fibrosis and inflammation, to capture genes implicated in EndoMT induction, regulation and phenotypic state across a range of physiological and pathological settings.

For each gene, the following annotations were recorded: HUGO Gene Nomenclature Committee (HGNC, RRID:SCR_002827) approved symbol and name (Seal *et al*., 2023), HGNC alias symbols and names, HGNC ID, HGNC gene group membership, Ensembl ID (release 114, RRID:SCR_002344) (Dyer *et al*., 2024), National Center for Biotechnology Information (NCBI) Entrez Gene ID (RRID:SCR_002473) (Maglott *et al*., 2005), and locus type including protein-coding and non-coding RNA classifications. Genes were manually assigned to one of seven marker subtype labels designed to capture endothelial identity, transitional cell states and EndoMT-inducing or regulatory activity associated with EndoMT transcriptional programming: canonical blood endothelial cell marker, canonical lymphatic endothelial cell marker, endothelial-associated marker, tumour endothelial marker, EndoMT inducer/inducer receptor, EndoMT positive regulator and mesenchymal-like cell marker. Definitions of each marker subtype are provided in Supplementary Data Sheet 1.

Genes were subsequently allocated to either the EndoMT UP or EndoMT DOWN gene set according to their reported or hypothesised direction of differential expression during partial EndoMT, defined here as an active transitional state rather than a complete mesenchymal endpoint with no expression of endothelial lineage genes. Partial EndoMT was selected as the reference state because many genes relevant to EndoMT biology represent context-dependent transitional-state responses, which may not be retained at a fully transitioned mesenchymal endpoint. The completed annotated curated gene set is provided in Supplementary Data Sheet 1.

### 2.2 EndoMT R Package

#### 2.2.1 Development and implementation of the EndoMT R package

The curated EndoMT gene sets were implemented as an openly available R package named EndoMT, available for download from GitHub (RRID:SCR_002630) at [https://github.com/kbdermdoc/EndoMT], with version 0.3.0 used in this study. The package was developed in R (RRID:SCR_001905) and requires R version 3.5.0 or later. The package is platform independent and is distributed under a GNU General Public License version 3.0.

The package contains three GMT files corresponding to the full EndoMT gene set and two directional subsets: ENDOTHELIAL_MESENCHYMAL_TRANSITION, ENDOTHELIAL_MESENCHYMAL_TRANSITION_UP and ENDOTHELIAL_MESENCHYMAL_TRANSITION_DOWN. Each GMT file is a tab-delimited plain-text file in which the first column contains the gene set name (string: free text), the second column contains a marker subtype descriptor (string: free text), and subsequent columns contain HGNC-approved gene symbols, with one gene identifier per field. Marker subtype descriptors correspond to the seven labels described in Section 2.1, in addition to the broader labels EndoMT, EndoMT UP and EndoMT DOWN, thereby enabling both broad EndoMT-focused GSEA and marker subtype-specific interrogation. The contents of each GMT file are provided in Supplementary Data Sheet 1.

The package includes three exported list objects: ENDOTHELIAL_MESENCHYMAL_TRANSITION, ENDOTHELIAL_MESENCHYMAL_TRANSITION_UP and ENDOTHELIAL_MESENCHYMAL_TRANSITION_DOWN, comprising the full EndoMT, EndoMT UP and EndoMT DOWN gene sets, respectively, together with their corresponding marker subtype-specific gene sets. The package also includes the annotation object EndoMT_geneset_info, which contains the curated gene-level metadata and marker subtype classifications described in Section 2.1. Package documentation is provided through R help pages for ENDOTHELIAL_MESENCHYMAL_TRANSITION and EndoMT_geneset_info, and incorporates example code for retrieving the GMT files.

#### 2.2.2 Package installation and gene set access

The EndoMT package can be installed from GitHub using devtools::install_github(“kbdermdoc/EndoMT”). The package can then be loaded using library(EndoMT). The full EndoMT gene set can be accessed using data(ENDOTHELIAL_MESENCHYMAL_TRANSITION) and the accompanying annotation data frame can be accessed using data(EndoMT_geneset_info).

The three packaged GMT files are stored in the package’s extdata directory and can be retrieved using system.file(). The GMT files can then be imported into R using clusterProfiler::read.gmt() (RRID:SCR_016884) for downstream GSEA. HGNC-approved gene symbols were checked against annotables (RRID:SCR_028420) reference tables corresponding to the Genome Reference Consortium Human GRCh37 and GRCh38 (RRID:SCR_006553) to confirm compatibility with both genome builds.

A complete installation and usage script illustrating installation of the EndoMT package, loading of the exported gene set and annotation objects, retrieval of the packaged GMT files for use in downstream GSEA, and verification of gene symbol compatibility with the GRCh37 and GRCh38 reference annotations, is provided in the accompanying GitHub repository.

### 2.3 Hallmark gene set overlap analysis

Human MSigDB (RRID:SCR_016863) collection H, Hallmark gene sets version 2025.1.Hs (Liberzon *et al*., 2015) were downloaded in April 2026 using RStudio (RRID:SCR_000432) with R version 4.5.1. The 50 Hallmark gene sets were exported as an Excel workbook (RRID:SCR_016137), including the corresponding gene set sizes (Supplementary Data Sheet 2). Gene overlap between the curated EndoMT gene set and each Hallmark gene set was then quantified by calculating the number of shared HGNC-approved gene symbols and the Jaccard index. Jaccard index was calculated as:

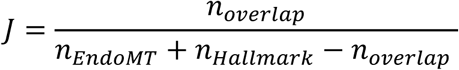

where *n*overlap represents the number of shared genes, and *n*EndoMT and *n*Hallmark represent the EndoMT and comparator Hallmark gene set sizes, respectively.

### 2.4 Benchmarking data sets

To assess the ability of the curated EndoMT gene sets to capture EndoMT-enriched biological processes, benchmarking was performed in four separate transcriptomic data sets (Table 1), first using the publicly available bulk RNA-seq data set GSE118446 (Caudrillier *et al*., 2020), in which EndoMT transcriptional activity was experimentally induced. In this data set, primary human umbilical vein endothelial cells (HUVECs) were treated *in vitro* for seven consecutive days with interleukin-1β (IL-1β; 1 ng/ml) alone, TGF-β2 (10 ng/ml) alone, or IL-1β and TGF-β2 combined (Monteiro *et al*., 2021). Human pulmonary arterial endothelial cells (HPAECs) were also co-treated. Treatments were performed in triplicate and compared with untreated control cells. RNA-seq was performed using ribosomal-depleted stranded libraries prepared by Beckman Coulter Genomics with the TruSeq Stranded Total RNA kit with Ribo-Zero Gold, followed by paired-end Illumina sequencing at an average depth of 50 million reads per sample (Monteiro *et al*., 2021).

**Table 1.** Transcriptomic data sets used for EndoMT gene set benchmarking analysis.

| **GEO Accession** | **DOI** | **Data type** | **Sample source / cells analysed** | **Treatment / comparison** |
| --- | --- | --- | --- | --- |
| GSE118446 | 10.1161/circresaha.120.318124 | Bulk RNA-seq | Primary ECs: HPAECs and HUVECs | IL-1β (1 ng/ml) and/or TGF-β2 (10 ng/ml) vs. Ctrl |
| GSE342578 | N/A | Bulk RNA-seq | Primary ECs: HUVECs | Combined TGF-β1/2/3 at 10, 2.5, 0.25 ng/ml vs. Ctrl |
| GSE237895 | 10.1126/sciadv.adp6182 | scRNA-seq | Primary ECs: HUVEC-derived mesenchymal-like ECs, tip and proliferating cells | TGF-β1 (10 ng/ml) or lentiviral SNAI1 transduction vs. Ctrl |
| GSE72056 | 10.1126/science.aad0501 | scRNA-seq | Human tissue: Melanoma metastases | CAFs vs. ECs |

CAFs, cancer-associated fibroblasts; Ctrl, control; ECs, endothelial cells; GEO, Gene Expression Omnibus; HPAECs, human pulmonary artery endothelial cell; HUVECs, human umbilical vein endothelial cell; N/A, not applicable; RNA-seq, RNA sequencing; scRNA-Seq, single-cell RNA sequencing.

Benchmarking was also performed using an in-house bulk RNA-seq data set GSE342578, in which HUVECs (PromoCell, C-12200) were cultured in Endothelial Cell Growth Medium (PromoCell, C-22010) at 37 °C in an atmosphere of 5% CO_2_, and were treated *in vitro* for three consecutive days with combined TGF-β1, TGF-β2 and TGF-β3 (PeproTech®, 17881453, 17860643 and 17831023, respectively, RRID:SCR_006802) at 0.25 ng/ml, 2.5 ng/ml or 10 ng/ml (defined here as low-, medium- and high-dose, respectively), prepared according to the manufacturer’s instructions. Treatments were performed in triplicate and compared with untreated control cells. Total cellular RNA was extracted using the Reliaprep™ RNA Miniprep System (Promega, Z6012) and sequenced by Azenta at a minimum sequencing depth of 20 million reads per sample. Sequencing reads were quality controlled, trimmed and aligned to the human genome using STAR (RRID:SCR_004463) with GENCODE v47 annotation (RRID:SCR_014966).

To demonstrate the utility of the EndoMT gene set in an alternative transcriptomic data type, benchmarking was also performed in two publicly available single-cell RNA sequencing (scRNA-seq) data sets. First, GSE237895 (Lebas *et al*., 2024) was used to assess two *in vitro* EndoMT models: HUVECs treated for seven consecutive days with TGF-β1 (10 ng/ml), and HUVECs with SNAI1 overexpression following overnight lentiviral SNAI1 transduction. These treatments were compared with untreated and pRRL-empty HUVEC controls, respectively. Single-cell droplet-based gene expression profiling was performed using libraries prepared with the Chromium Single Cell 3′ Library, Gel Bead & Multiplex Kit, and Chip Kit (10x Genomics, v3.1), followed by MGISEQ-2000 sequencing (Lebas *et al*., 2024). Second, GSE72056 was used as a human tumour-derived data set comprising 4645 single cells isolated from 19 freshly procured human melanoma samples, including lymphoid metastases (n = 10), distant organ metastases (n = 8) and one primary acral melanoma. Samples were disaggregated and underwent Smart-Seq2 library preparation, followed by Illumina NextSeq 500 sequencing (Tirosh *et al*., 2016).

#### 2.4.1 Analysis of GSE118446, GSE342578 and GSE237895

##### 2.4.1.1 Differential expression analysis

RNA-seq raw counts matrices were downloaded from Gene Expression Omnibus (GEO, RRID:SCR_005012) (Edgar, Domrachev and Lash, 2002) accessions GSE118446 and GSE342578 using R. Sample metadata were extracted and aligned with the count matrix using GEO Sample (GSM) identifiers. Differential expression analysis was performed using DESeq2 (RRID:SCR_015687) (Love, Huber and Anders, 2014) separately for each cell type and treatment contrast to generate contrast-specific differential expression results for benchmarking. For each contrast, samples were subset to the relevant cell type and treatment comparison. For GSE118446 only, low-count genes were pre-filtered separately for each contrast using a total raw count threshold of at least 10 (Love, Anders and Huber, 2023). For GSE342578, no additional low-count filtering was applied. For both data sets, differential expression was modelled using a design formula of ∼treatment, with the control group specified as the reference level (Supplementary Data Sheets 3-4).

For GSE237895, previously generated differential expression results provided as supplementary material by Lebas et al. were used (Lebas *et al*., 2024). Lebas et al. performed pairwise differential analysis comparing treatment and control conditions within each endothelial cell subpopulation using limma, and the resulting supplementary differential expression tables were used for benchmarking (Haghverdi *et al*., 2018; Lebas *et al*., 2024).

##### 2.4.1.2 Gene Set Enrichment Analysis

TERM2GENE files were generated for the Hallmark, EndoMT UP and EndoMT DOWN gene sets (Supplementary Data Sheet 5). Hallmark gene sets were retrieved using msigdbr (RRID: SCR_022870), while EndoMT gene sets were retrieved from the EndoMT R package as described in Section 2.2.2. Gene identifiers were standardised to NCBI Entrez Gene IDs for compatibility with downstream GSEA.

Ranked gene lists were generated from the differential expression results described above, with genes ranked in decreasing order by log2 fold change (log2FC) to be used as input for GSEA (RRID:SCR_003199) (Subramanian *et al*., 2005). GSEA was performed using clusterProfiler (RRID:SCR_016884) (Yu *et al*., 2012) across all Hallmark gene sets and the curated EndoMT UP and EndoMT DOWN gene sets, with or without inclusion of the seven marker subtype-specific gene sets described in Section 2.1. A weighting exponent of 1 was used, with minimum and maximum gene set size thresholds of 4 and 500 genes, respectively. The nominal *P* value cutoff was set to 1 to retain all enrichment results for downstream comparison, and *P* values were adjusted using the Benjamini-Hochberg (BH) method. GSEA outputs were saved as .rds files.

##### 2.4.1.3 Integrated EndoMT and Hallmark Gene Set Enrichment Analysis visualisation

For each data set, integrated GSEA result objects were generated for each contrast. Hallmark, EndoMT UP and EndoMT DOWN GSEA results were combined using dplyr::bind_rows() (RRID:SCR_016708). For marker subtype-level analyses, Hallmark results were similarly combined with EndoMT marker subtype GSEA results. A contrast identifier was appended to each combined data frame and the resulting contrast-level integrated result objects were saved as .rds files; these are available in the accompanying GitHub repository.

Integrated GSEA results were visualised using GeneRatio dot plots and normalised enrichment score (NES) bar plots generated with ggplot2 (RRID:SCR_014601). A complete worked example, including the code used to generate these visualisations, is available for the GSE118446 data set in the accompanying GitHub repository.

##### 2.4.1.4 Leading edge subset Gene Ontology enrichment analysis

Leading edge genes were extracted from the core_enrichment column of GSEA result files across all contrasts. To interpret the biological processes represented among genes driving significant GSEA signals, Gene Ontology (GO, RRID:SCR_002811) (Blake *et al*., 2013) Biological Process enrichment analysis was performed on leading edge gene subsets derived from significantly enriched GSEA terms, defined by a BH-adjusted *P* value ≤ 0.05. Leading edge gene subsets containing fewer than ten genes were excluded. GO Biological Process enrichment was performed using clusterProfiler::enrichGO() with org.Hs.eg.db, keyType = “ENTREZID”, ont = “BP”, and BH *P* value adjustment, *P* value and q-value cutoffs of ≤ 0.05. For each contrast, the background universe was defined as the ranked gene list used as input for the corresponding GSEA. Full unsimplified GO Biological Process enrichment results are available in Supplementary Data Sheet 6. For data visualisation, enriched GO terms were simplified using clusterProfiler::simplify() with a semantic similarity cutoff of 0.7 to reduce redundancy among related GO terms.

#### 2.4.2 Analysis of GSE72056

The RNA-seq count matrix was downloaded from GEO was analysed using Seurat version 5 (RRID:SCR_007322). Cell populations were identified according to their original cell-type annotations provided by Tirosh et al. (Tirosh *et al*., 2016). Endothelial cells (n = 65) and CAFs (n = 61) were included in the analysis, to provide a human-derived tissue model of EndoMT gene set enrichment between vascular and stromal populations. For each gene, log2FC values were calculated by comparing mean expression in CAFs versus endothelial cells. Positive log2FC values indicated higher expression in CAFs and negative log2FC values indicated higher expression in endothelial cells. Genes were ranked by log2FC and used as input for GSEA with the EndoMT UP and EndoMT gene sets.

## 3 Results

### 3.1 EndoMT is distinct from existing Hallmark gene sets

A bespoke 193-gene EndoMT signature was curated through comprehensive review of the EndoMT literature, comprising 140 genes anticipated to be upregulated, and 53 genes anticipated to be downregulated during this dynamic transitional process. To determine whether the curated EndoMT gene signature captured biology distinct from existing MSigDB Hallmark gene sets (Liberzon *et al*., 2015), overlap analysis was performed against all 50 Hallmark signatures. The EndoMT gene set showed the greatest overlap with the 200-gene Hallmark EMT signature (40 overlapping genes; Jaccard index = 0.11; Figure 1), followed by the 200-gene Hallmark apical junction signature (24 overlapping genes; Jaccard index = 0.07; Figure 1). Overlap with all other Hallmark gene sets was minimal (Supplementary Figure 1, Supplementary Data Sheet 2), supporting the EndoMT gene set as a distinct resource for transcriptomic enrichment analysis that complements existing Hallmark pathway signatures.

**Figure 1.**
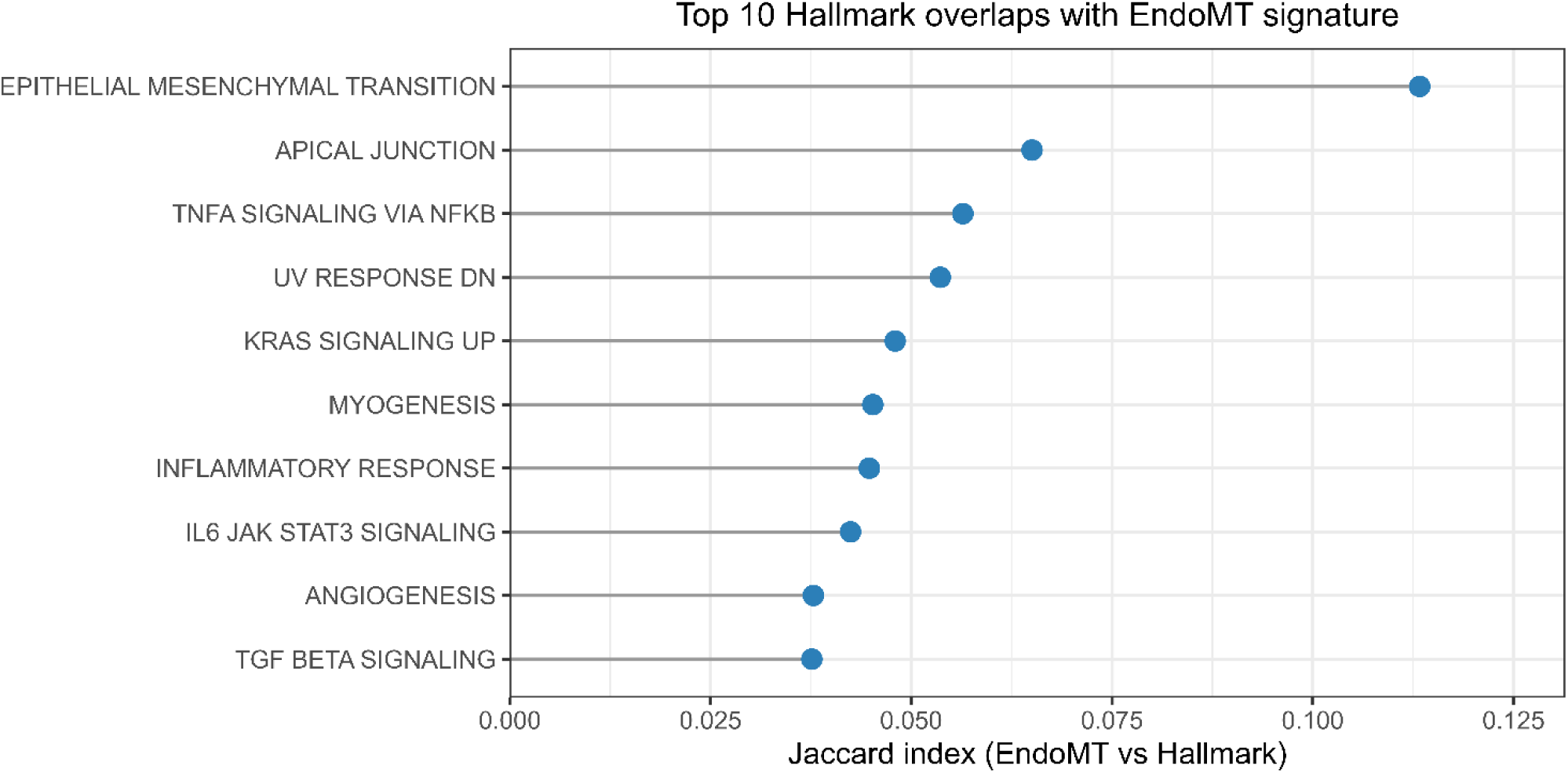
EndoMT overlaps most strongly with Hallmark EMT. Lollipop chart showing the top ten MSigDB Hallmark gene set overlaps with the curated EndoMT signature, ranked by Jaccard index.

### 3.2 TGF-β-associated EndoMT signatures are context-, dose- and duration-dependent in primary human endothelial cells

#### 3.2.1 EndoMT gene sets confirm IL-1β/TGF-β2 co-treatment induces EndoMT in GSE118446 HUVECs and HPAECs

The publicly available bulk RNA-seq data set GSE118446 was analysed to benchmark the curated EndoMT gene sets against an established EndoMT-positive transcriptional data set. This included HUVECs and HPAECs treated with IL-1β and TGF-β2, either alone or in combination, for seven days. In the original study, real-time quantitative polymerase chain reaction (RT-qPCR) analysis confirmed induction of EndoMT in co-treated cells, with loss of endothelial cell marker Platelet And Endothelial Cell Adhesion Molecule 1 (PECAM1), gain of mesenchymal cell markers Actin Alpha 2 Smooth Muscle (ACTA2) and Collagen Type I Alpha 1 Chain (COL1A1), and expression of EndoMT positive regulator Snail Family Transcriptional Repressor 2 (SNAI2) (Monteiro *et al*., 2021). Changes in PECAM1 and SNAI2 expression were further validated at the protein level by immunofluorescence microscopy, while reduced PECAM1 and increased ACTA2 were also confirmed by Western blot in HPAECs. Consistent with these reported findings, GSEA of the co-treated HUVEC and HPAEC contrasts demonstrated significant positive enrichment of the EndoMT UP gene set and significant negative enrichment of the EndoMT DOWN gene set (Figure 2A-D).

**Figure 2.**
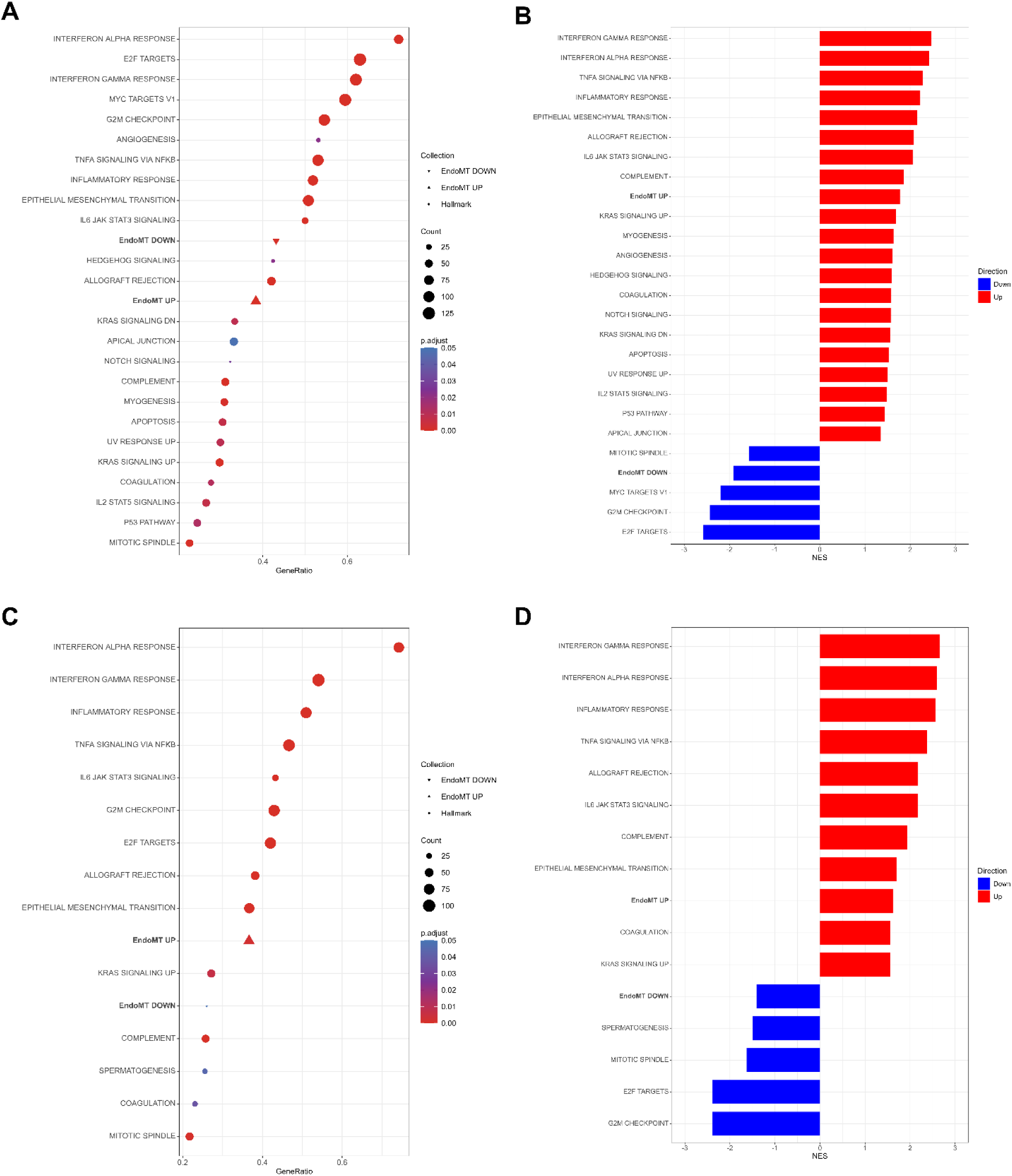
Application of EndoMT R package gene sets confirms EndoMT enrichment following IL-1β/TGF-β2 co-treatment in GSE118446 endothelial cells. GSEA was performed on GSE118446 HUVECs and HPAECs treated with combined IL-1β/TGF-β2 for seven days compared with untreated controls, using EndoMT R package-derived EndoMT UP and EndoMT DOWN gene sets, alongside MSigDB Hallmark gene sets. GSEA result objects were integrated prior to visualisation. Gene ratio dot plots show significantly enriched gene sets in (A) HUVECs and (B) HPAECs. NES bar plots show the direction and magnitude of enrichment in (C) HUVECs and (D) HPAECs. Only gene sets passing BH-adjusted *P* value ≤ 0.05 are shown.

The GMT files were additionally structured to permit exploratory GSEA at the level of curated EndoMT marker subtypes, as defined in Section 2.1 and Supplementary Data Sheet 1. In these analyses, broad EndoMT UP or EndoMT DOWN terms were tested alongside their respective marker subtype terms. Consistent with acquisition of an EndoMT phenotype, the mesenchymal-like cell marker subtype showed significant positive enrichment in both co-treated HUVECs and HPAECs (Supplementary Figure 2A-B). Overall, broad EndoMT UP and EndoMT DOWN enrichment signals were retained across the marker subtype GSEA outputs, with the exception of EndoMT DOWN in co-treated HPAECs, which was significant in the broad EndoMT DOWN analysis (BH-adjusted *P* value = 0.0488) but not in the marker subtype analysis (BH-adjusted *P* value = 0.1084) (Supplementary Figure 2B). Given the increased number and altered composition of tested gene sets, together with the small size of several subtype-specific gene sets, marker subtype-level enrichment was interpreted as supportive benchmarking evidence rather than as a primary enrichment readout.

Leading edge inspection further supported concordance with the original RT-qPCR findings. In HUVECs, the EndoMT UP leading edge included *COL1A1* and *SNAI2*, while the EndoMT DOWN leading edge included *PECAM1*. In HPAECs, the EndoMT UP leading edge included *ACTA2*. Beyond these directly overlapping markers, shared EndoMT UP leading edge genes across HUVECs and HPAECs captured broader EndoMT-associated biology, including EndoMT-inducing ligands and signalling mediators such as *IL1A, IL1B, IL6, TNF, TGFB2, TGFBR1, NFKB1, NFKB2* and *RELB*; transcriptional regulators *SNAI1* and *HEY1*; mesenchymal cell markers, extracellular matrix and remodelling genes such as *TNC*, *TGFBI, TAGLN, CNN1, MYL9, LAMC2, COL4A1, COL4A2, FN1, FAP, ADAM12, SERPINE1* and *CDH2*, in addition to inflammatory adhesion molecules *VCAM1* and *ICAM1* (Figure 3A, C). Conversely, shared EndoMT DOWN leading edges included endothelial lineage, junctional and vascular homeostasis genes such as *CLDN5, ICAM2, VWF, EMCN, PROX1, LYVE1, MIR126, ACE, GJA4, NECTIN1* and *ADAMTS18* (Figure 3B, D). Full ranked leading edge gene lists are provided in Supplementary Data Sheet 7.

**Figure 3.**
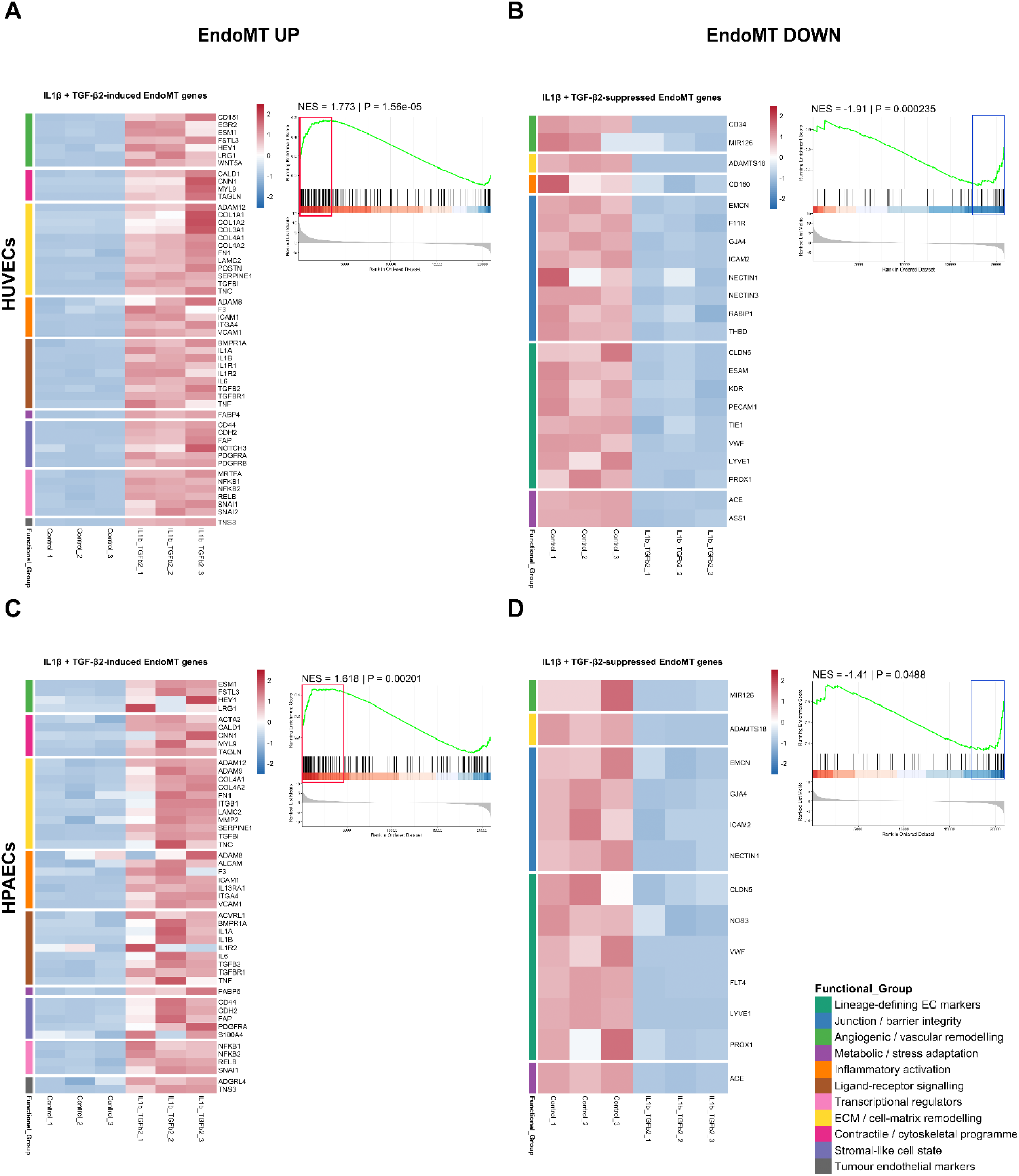
Leading edge analysis of EndoMT UP and EndoMT DOWN gene set enrichment in GSE118446 IL-1β/TGF-β2 co-treated HUVECs and HPAECs. Enrichment plots and corresponding heatmaps of the leading edge genes grouped by functional annotation for (A, B) HUVECs and (C, D) HPAECs.

Benchmarking analyses demonstrate that the EndoMT UP and EndoMT DOWN gene sets detect transcriptional changes consistent with experimentally validated EndoMT induction in GSE118446 HUVECs and HPAECs, particularly following combined IL-1β/TGF-β2 stimulation.

#### 3.2.2 EndoMT gene sets reveal partial activation of EndoMT-associated transcriptional programmes in IL-1β or TGF-β2-treated GSE118446 HUVECs

Although the original RT-qPCR and phenotypic analyses did not show IL-1β or TGF-β2 treatment alone induced overt EndoMT (Monteiro *et al*., 2021), GSEA of bulk RNA-seq from IL-1β- or TGF-β2-treated HUVECs in GSE118446 demonstrated significant positive enrichment of the EndoMT UP gene set and significant negative enrichment of the EndoMT DOWN gene set (Figure 4A-D). These findings suggest that single ligand stimulation activates components of the EndoMT transcriptional programme in HUVECs, without necessarily recapitulating a more coordinated marker-defined EndoMT phenotype observed following combined IL-1β/TGF-β2 treatment.

**Figure 4.**
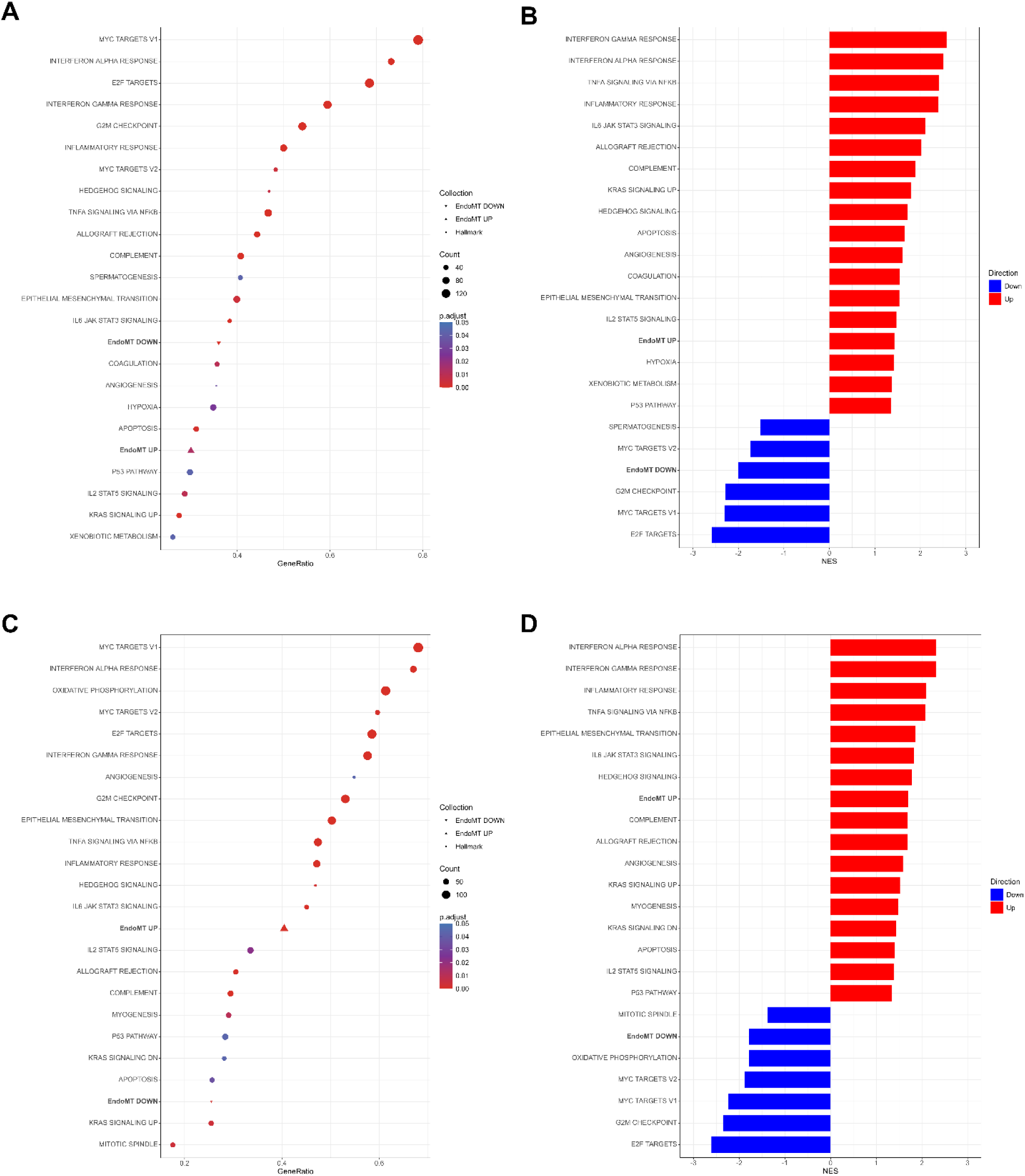
EndoMT R package gene sets detect directional EndoMT-associated enrichment following single ligand treatment in GSE118446 HUVECs. GSEA was performed on GSE118446 HUVECs treated with IL-1β or TGF-β2 for seven days compared with untreated controls, using EndoMT R package-derived EndoMT UP and EndoMT DOWN gene sets, alongside MSigDB Hallmark gene sets. Gene ratio dot plots and corresponding NES bar plots show significantly enriched gene sets in (A, B) IL-1β-treated HUVECs and (C, D) TGF-β2-treated HUVECs. Only gene sets passing BH-adjusted *P* value ≤ 0.05 are shown.

Leading edge analysis suggested that each single treatment captured distinct components of the EndoMT programme. In IL-1β-treated HUVECs, the EndoMT UP leading edge was dominated by inflammatory and activation-associated genes, including EndoMT-inducing ligands *IL1A*, *IL1B* and *IL6*; NF-κB signalling mediators and transcriptional regulators *NFKB1*, *NFKB2* and *RELB*; inflammatory adhesion molecules *VCAM1* and *ICAM1*; and mesenchymal cell markers, extracellular matrix and remodelling genes such as *TNC*, *ACTA2*, *TAGLN*, *MYL9*, *COL4A2*, *FAP*, *ADAM12* and *SERPINE1* (Supplementary Figure 3A). Conversely, the EndoMT DOWN leading edge included endothelial lineage and vascular homeostasis genes such as *PECAM1*, *ICAM2*, *VWF*, *EMCN*, *NOS3*, *PROX1*, *LYVE1*, *FLT4*, *CD34*, *ACE*, *GJA4* and *GJA5* (Supplementary Figure 3B).

In TGF-β2-treated HUVECs, the EndoMT UP leading edge showed a more matrix-remodelling and mesenchymal-regulatory profile, including EndoMT-inducing ligands and signalling mediators such as *IL1A, IL1R2, IL6, TGFB2*, *TGFBR1* and *TNF*; transcriptional regulators *SNAI1*, *SNAI2, NFKB1* and *NFKB2*; and mesenchymal cell markers, extracellular matrix and remodelling genes such as *COL1A1*, *COL1A2*, *COL3A1*, *TNC*, *TAGLN*, *CNN1*, *MYL9*, *FN1*, *POSTN*, *FAP*, *ADAM12*, *MMP2* and *SERPINE1* (Supplementary Figure 3C). Conversely, the EndoMT DOWN leading edge included endothelial lineage and vascular homeostasis genes such as *MIR126*, *ACKR1*, *VWF*, *ICAM2*, *CLDN5*, *ACE*, *GJA4*, *CD34* and *LYVE1* (Supplementary Figure 3D).

Together, these analyses show that the EndoMT gene sets capture broader transcriptional changes consistent with an early or partial EndoMT-associated activation in single ligand-treated HUVECs. These findings support the utility of the EndoMT gene sets for identifying intermediary EndoMT-associated transcriptional states that may not otherwise be fully resolved by a focused marker panel alone.

#### 3.2.3 EndoMT gene sets capture divergent enrichment following early versus chronic TGF-β exposure in HUVECs

To assess whether the curated EndoMT gene sets distinguished early from more established TGF-β-associated EndoMT transcriptional responses, in-house bulk RNA-seq data from GSE342578 HUVECs treated with combined TGF-β1/2/3 for three days were compared with seven-day ligand-treated GSE118446 HUVECs. GSEA showed significant negative enrichment of the EndoMT UP gene set in high-dose TGF-β1/2/3-treated HUVECs, in contrast to the positive enrichment observed across IL-1β/TGF-β2-, IL-1β- or TGF-β2-treated GSE118446 HUVECs. The EndoMT DOWN gene set was not significantly enriched in the high-dose treatment arm, and neither gene set was significantly enriched in the medium- or low-dose treatment arms (Supplementary Figure 4A-B). Additional exploratory analysis at the level of the curated EndoMT marker subtypes, as defined in Section 2.1 and Supplementary Data Sheet 1, further supported this divergent enrichment pattern. High-dose TGF-β1/2/3 treatment showed significant negative enrichment of EndoMT positive transcriptional regulators, mesenchymal-like cell markers and tumour endothelial markers (Figure 5A). The overall enrichment profile suggests that early high-dose TGF-β1/2/3 exposure did not induce early, partial or established EndoMT-associated transcriptional activity.

**Figure 5.**
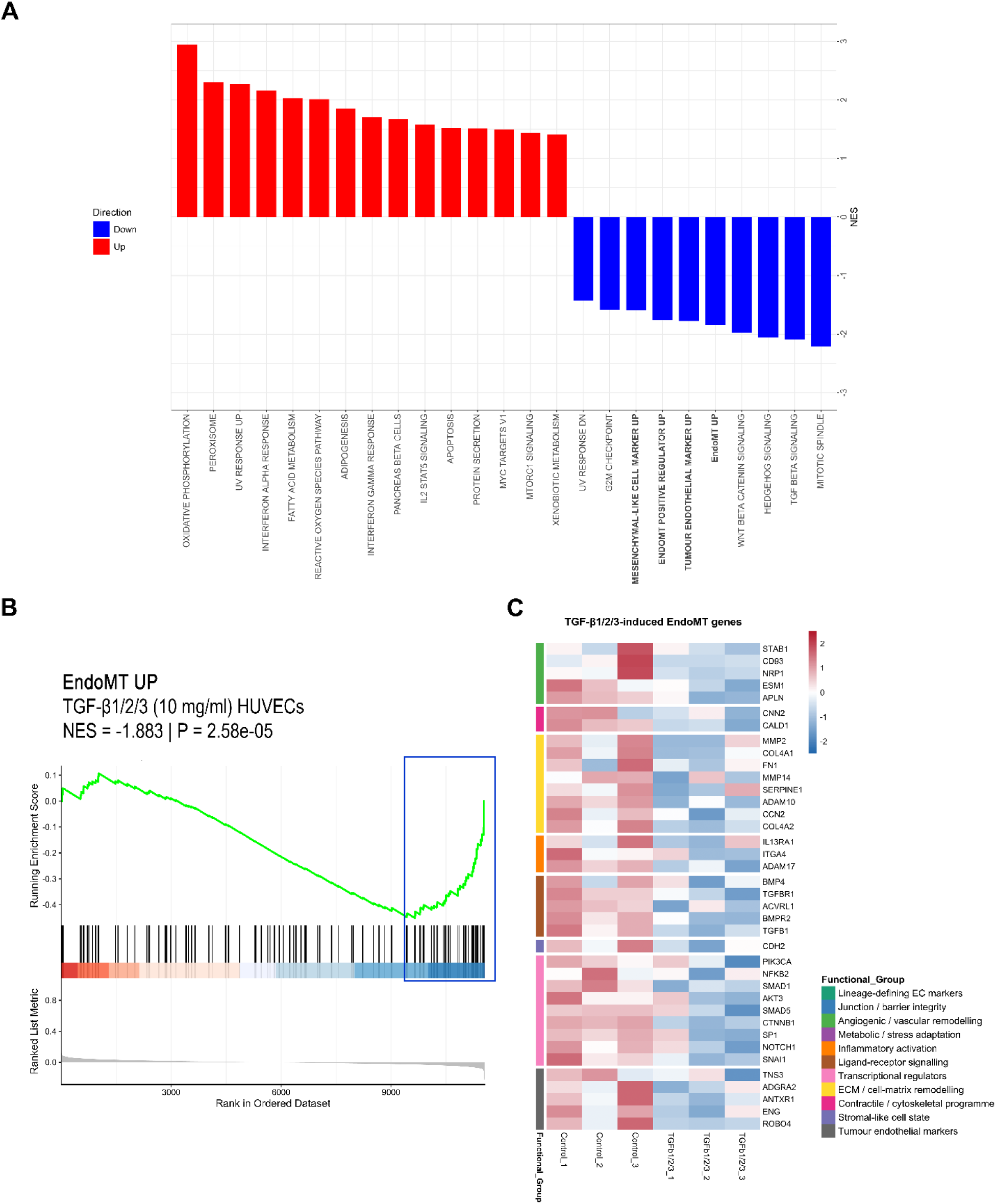
Application of EndoMT R package gene sets identifies negative enrichment of EndoMT-associated signatures following early high-dose TGF-β1/2/3 exposure in HUVECs. GSEA was performed on in-house GSE342578 HUVECs treated with combined high-dose TGF-β1/2/3 for three days compared with untreated controls, using EndoMT R package-derived gene sets, alongside MSigDB Hallmarks. (A) NES bar plot of EndoMT UP and EndoMT DOWN marker subtype-specific EndoMT gene set enrichment, showing significant negative enrichment of EndoMT UP signature, EndoMT positive transcriptional regulators, mesenchymal-like cell markers and tumour endothelial markers. EndoMT DOWN was not significantly enriched. Only gene sets passing BH-adjusted *P* value ≤ 0.05 are shown. (B) Enrichment plot of the negatively enriched EndoMT UP gene set. (C) Heatmap of the EndoMT UP leading edge genes grouped by functional annotation.

Leading edge inspection of the negatively enriched EndoMT UP gene set included TGF-β/bone morphogenetic protein (BMP), developmental and transcriptional signalling mediators such as *TGFB1*, *BMP4*, *TGFBR1*, *ACVRL1*, *SMAD1*, *SMAD5*, *PIK3CA*, *AKT3*, *CTNNB1*, *NFKB2*, *NOTCH1* and *SNAI1*; mesenchymal cell markers, extracellular matrix and remodelling genes including *FN1*, *COL4A1*, *COL4A2*, *MMP2*, *MMP14*, *SERPINE1*, *ADAM10*, *ADAM17*, *TNS3*, *CDH2*, *CALD1*, *CNN2* and *CCN2*, in addition to adhesion-associated *ITGA4* and tumour endothelial marker *ROBO4* (Figure 5B-C). Notably, several leading edge genes overlapped with those contributing to positive EndoMT UP enrichment in ligand-treated GSE118446 HUVECs, suggesting that the divergent enrichment reflected differential regulation of a shared subset of EndoMT-associated genes. Full ranked leading edge gene lists are provided in Supplementary Data Sheet 8.

These findings indicate that three-day high-dose TGF-β1/2/3 exposure induced a transcriptional state distinct from the positive EndoMT programme observed after chronic ligand exposure in GSE118446 HUVECs, characterised by relative repression of EndoMT-associated genes without significant evidence of endothelial lineage loss.

### 3.3 EndoMT UP gene set captures vascular-mesenchymal biology beyond Hallmark EMT

GO Biological Process enrichment analysis was performed on leading edge genes derived from EndoMT UP and Hallmark EMT gene sets with significant positive enrichment in GSE118446, to evaluate the biological processes driving enrichment and to assess whether the EndoMT UP gene set captured mesenchymal biology distinct from the existing Hallmark EMT signature. In IL-1β/TGF-β2 co-treated HUVECs and HPAECs, Hallmark EMT leading edge genes enriched broad stromal, extracellular matrix, wound-response, adhesion and developmental processes, including “extracellular matrix organization”, “response to wounding”, “positive regulation of cell adhesion”, “cell-matrix adhesion”, “cell-substrate adhesion” and endoderm-associated developmental terms. EndoMT UP leading edge genes showed overlap with several of these EMT-associated processes, but also enriched additional vascular-remodelling, migratory and mesenchymal-related processes; these included “positive regulation of locomotion”, “mesenchymal cell differentiation” and “mesenchyme development”, as well as inflammation-associated EndoMT processes such as “acute inflammatory response”, “leukocyte migration”, and “leukocyte adhesion to vascular endothelial cell”. Where shared vascular or migratory terms were observed, including “angiogenesis” and “positive regulation of cell migration”, these terms showed higher gene ratios and lower adjusted *P* values in the EndoMT UP leading edge analysis than in the corresponding Hallmark EMT analysis (Figure 6A-D).

**Figure 6.**
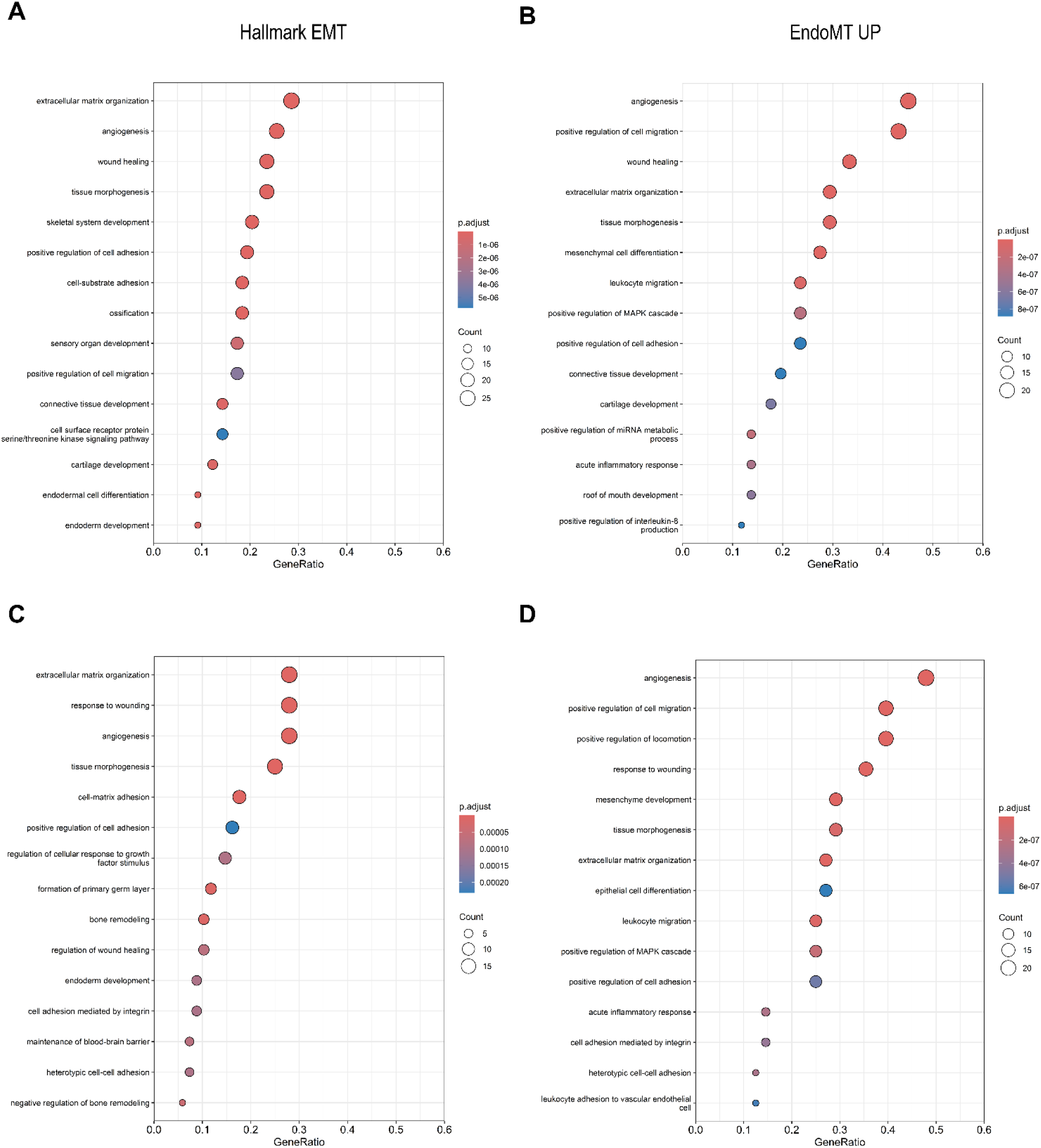
GO Biological Process enrichment of Hallmark EMT and EndoMT R package-derived EndoMT UP leading edge genes in GSE118446 co-treated endothelial cells. GO Biological Process enrichment analysis was performed on leading edge gene subsets derived from significantly positively enriched Hallmark EMT and EndoMT UP gene sets in GSE118446 HUVECs and HPAECs treated with combined IL-1β/TGF-β2 for seven days compared with untreated controls. Enrichment plots are shown for Hallmark EMT and EndoMT UP leading edge genes in (A, B) HUVECs and (C, D) HPAECs. GO terms were filtered by BH-adjusted *P* value ≤ 0.05.

Similar vascular, mesenchymal and inflammatory biological processes were observed in single ligand-treated HUVECs (Supplementary Figure 5A-D), where the EndoMT UP gene set showed significant positive enrichment (Figure 4B, D) despite overt EndoMT induction not being validated in the original study by Monteiro et al. (Monteiro *et al*., 2021). This supports the EndoMT UP gene set as an EMT-related but distinct signature capable of enriching for vascular-mesenchymal biological processes relevant to EndoMT.

### 3.4 EndoMT UP gene set detects TGF-β1 and SNAI1-associated EndoMT programmes in GSE237895 single-cell endothelial subpopulations

The publicly available scRNA-seq data set GSE237895 was analysed to benchmark the curated EndoMT gene sets in an alternative transcriptomic data type and across transdifferentiating endothelial cell subpopulations in established EndoMT models. GSE237895 HUVECs treated with TGF-β1 for seven days showed induction of mesenchymal markers with concurrent reduction in endothelial identity markers. In a separate model, GSE237895 HUVECs underwent SNAI1 overexpression following lentiviral SNAI1 transduction, with functional consequences validated by *in vitro* assays demonstrating increased migration and reduced transendothelial electrical resistance, consistent with impaired endothelial barrier integrity and EndoMT induction (Lebas *et al*., 2024). Single-cell profiling identified endothelial-derived subpopulations across these models, including mesenchymal-like endothelial cells characterised by expression of SERPINE1 and FN1, proliferating endothelial cells defined by MKI67 and PTTG1 expression, and angiogenic tip cells defined by PGF and CXCR4 expression (Lebas *et al*., 2024).

Consistent with an activated EndoMT transcriptional programme, GSEA of the mesenchymal-like endothelial cells from both TGF-β1-treated and SNAI1-overexpressing HUVECs demonstrated significant positive enrichment of the EndoMT UP gene set, with non-significant negative enrichment of the EndoMT DOWN gene set (Supplementary Figure 6A). Leading edge inspection showed that the shared EndoMT UP signal across both models was dominated by mesenchymal cell markers, extracellular matrix and remodelling genes, including *FN1*, *CCN2*, *MMP2*, *SPARC*, *COL4A1*, *ITGB1*, *SERPINE1*, *COL4A2*, *COL3A1*, *CALD1*, *CNN3*, *ADAM9* and *ADAM10*; EndoMT-associated signalling mediators *BMP4* and *TGFBR2*; adhesion and inflammatory-associated genes *ITGA4* and *ICAM1*; and tumour endothelial marker genes *ENG* and *BGN*. These leading edge profiles support concordance between EndoMT UP enrichment and mesenchymal transdifferentiation-associated endothelial cell states across both chronic TGF-β1-stimulation and SNAI1-overexpression models.

In TGF-β1-treated tip cells, GSEA also demonstrated significant positive enrichment of the EndoMT UP gene set, while EndoMT DOWN showed non-significant positive enrichment (Supplementary Figure 6B). The EndoMT UP leading edge in this subpopulation included angiogenic, endothelial-associated and tumour endothelial marker genes such as *APLN*, *ENG*, *ROBO4*, *CLEC14A*, *NRP1*, *ESM1*, *ADGRA2*, *PLVAP, STAB1* and *BGN*; mesenchymal cell markers, extracellular matrix and remodelling genes, including *CCN2*, *SERPINE1*, *MMP2*, *COL4A1*, *COL4A2*, *FN1*, *SPARC*, *CALD1* and *CNN3*; EndoMT-associated signalling mediators *BMP4*, *CTNNB1*, *AKT3* and *FOXC2*. The positively enriched EndoMT DOWN leading edge retained endothelial lineage and vascular homeostasis genes including *VWF*, *PROCR*, *ERG*, *FLT1*, *FGFR1*, *ACE*, *FLT4*, *CD34*, *PODXL* and *ECSCR*. These enrichment patterns are consistent with a partial EndoMT-associated tip cell phenotype observed during sprouting angiogenesis, in which migratory and vascular-remodelling programmes are activated while endothelial identity is retained (Jakobsson *et al*., 2010; Welch-Reardon, Wu and Hughes, 2015).

In proliferating TGF-β1-treated HUVECs, GSEA demonstrated significant positive enrichment of the EndoMT UP gene set, together with significant positive enrichment of the EndoMT DOWN gene set (Supplementary Figure 6C). The EndoMT UP leading edge in this subpopulation included mesenchymal cell markers, extracellular matrix, adhesion and remodelling genes such as *SERPINE1*, *COL4A1*, *COL4A2*, *MMP2*, *FN1*, *CCN2*, *SPARC*, *CNN3*, *CALD1*, *CDH2*, and *ADAM17*; EndoMT-inducing ligands and signalling mediators, including *BMP4*, *TGFB1*, *ACVRL1* and *TNFRSF1B*; transcriptional regulators *CTNNB1*, *NOTCH1*, *FOXC2*, *SMAD1* and *SMAD4*; angiogenic, endothelial-associated and tumour endothelial marker genes, including *ENG*, *ROBO4*, *APLN*, *CLEC14A*, *ADGRA2*, *PLVAP*, *ESM1*, *STAB1*, *EGFL7*, *KIT* and *BGN*. The inclusion of *TGFB1*, *ACVRL1* and *SMAD1* is consistent with transcriptional representation of the TGF-β1/ALK1/SMAD signalling axis implicated in endothelial proliferation and migration (Goumans *et al*., 2003), while also situating this leading edge within broader TGF-β/BMP signalling biology relevant to endothelial plasticity and EndoMT (Yoshimatsu and Watabe, 2022; Kovacic *et al*., 2019). The positively enriched EndoMT DOWN leading edge included endothelial lineage and vascular homeostasis genes such as *VWF*, *TIE1*, *FLT1*, *CDH5*, *PECAM1*, *TEK*, *ERG* and *KDR*. These enrichment patterns support a partial EndoMT-associated phenotype in proliferating endothelial cells, characterised by matrix-remodelling and TGF-β-associated programmes, alongside retention of endothelial identity. Full ranked leading edge gene lists are provided in Supplementary Data Sheet 9.

Overall, benchmarking in GSE237895 demonstrated that the EndoMT UP gene set detects EndoMT-associated transcriptional programmes in scRNA-seq-derived endothelial cell subpopulations, particularly in the mesenchymal-like endothelial cells from chronic TGF-β1- and SNAI1-overexpression models. In contrast, EndoMT DOWN enrichment was more heterogeneous across subpopulations, supporting interpretation of EndoMT as a context-dependent and sometimes partial endothelial cell state-transition.

### 3.5 EndoMT gene sets discriminate mesenchymal and endothelial states in human melanoma tissue

The publicly available melanoma scRNA-seq data set GSE72056 was used to assess the curated EndoMT gene sets in human tissue, in contrast to the *in vitro* models discussed. Using a CAF versus endothelial cell contrast, GSEA demonstrated significant positive enrichment of the EndoMT UP gene set and significant negative enrichment of the EndoMT DOWN gene set in CAFs compared with endothelial cells (Supplementary Figure 7A-B). This pattern is consistent with the mesenchymal and matrix-remodelling phenotype of CAFs and is compatible with evidence that EndoMT is an established source of CAF populations in the tumour microenvironment (Zeisberg *et al*., 2007a; Potenta, Zeisberg and Kalluri, 2008).

Inspection of the most differentially expressed genes further supported this finding. Among the genes with the highest positive log2FC in CAFs versus endothelial cells were several EndoMT UP genes, including mesenchymal-like cell markers *COL1A2*, *COL1A1*, *COL3A1*, *PCOLCE*, *TGFBI*, *S100A4, CD44, TAGLN, ACTA2, PDGFRB* and *MYL9*, tumour endothelial markers *CD248* and *ANTXR1*, and EndoMT regulator *SNAI2* (Figure 7). Complementary to this, among the genes with the most negative log2FC were canonical blood endothelial markers *CLDN5*, *VWF*, *CDH5*, *PECAM1*, *ERG* and *ESAM*, and canonical lymphatic endothelial markers *LYVE1* and *FLT4* from the EndoMT DOWN gene set. Together these results support the ability of the EndoMT UP and EndoMT DOWN gene sets to discriminate mesenchymal and endothelial transcriptional states within human melanoma tissue. A complete list of differentially expressed genes is available in Supplementary Data Sheet 10.

**Figure 7.**
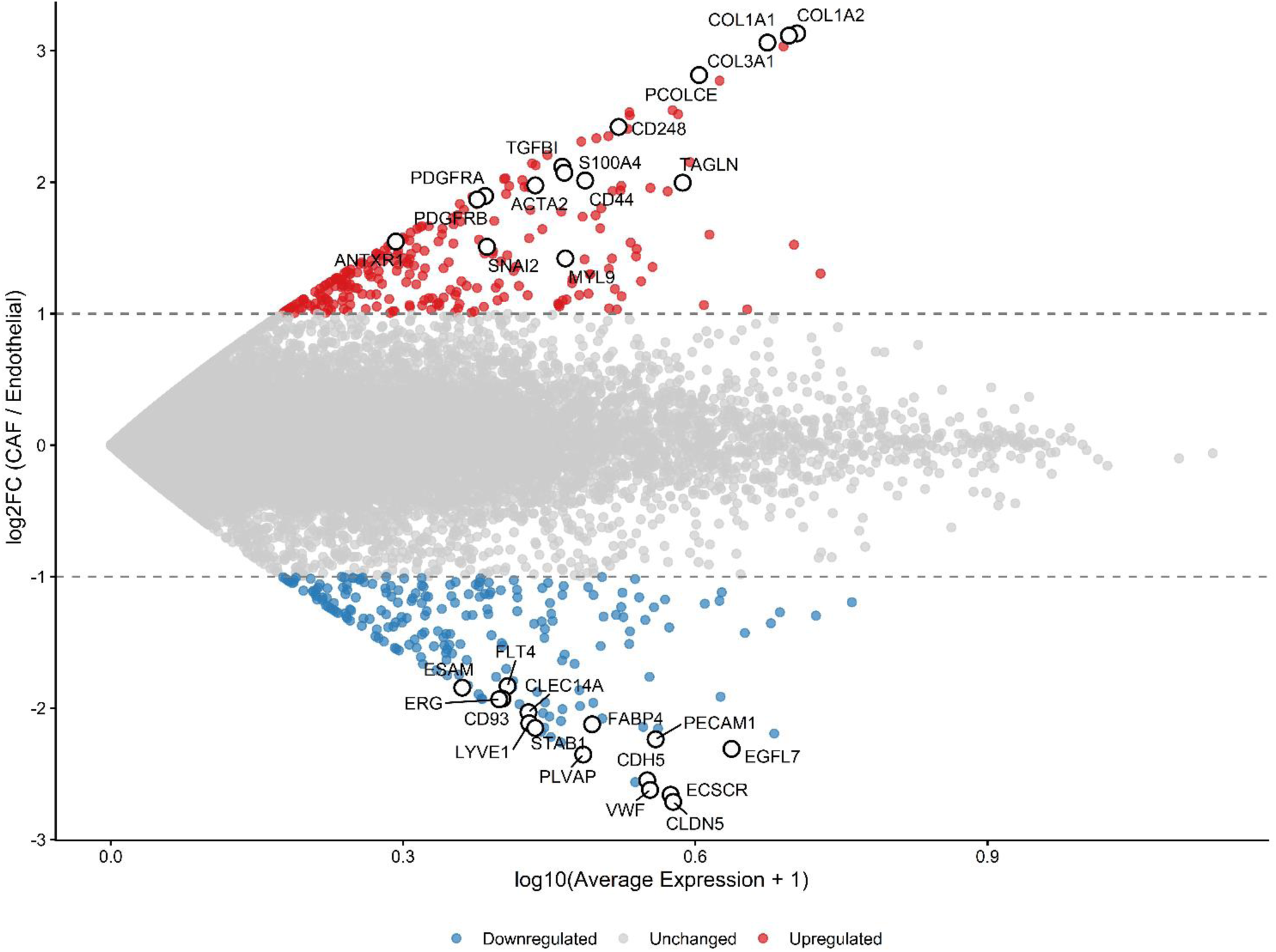
EndoMT gene sets discriminate mesenchymal and endothelial transcriptional states in human melanoma tissue. Mean-average (MA) plot of GSE72056 scRNA-seq data comparing CAFs with endothelial cells. For each gene, average expression was calculated within each cell population and used to derive log2FC in CAFs versus endothelial cells. Annotated genes represent the fifteen most positively and negatively differentially expressed genes in CAFs versus endothelial cells.

## 4 Discussion

In this study, we present a curated EndoMT gene set resource and accompanying R package designed to support transcriptomic analysis of EndoMT across experiment and disease contexts. The resource provides directionally assigned EndoMT UP and EndoMT DOWN gene sets, marker subtype annotations, GMT files and exported R objects, with example workflows for integration with commonly used GSEA pipelines available in the accompanying GitHub repository. Benchmarking analyses using publicly available and in-house bulk RNA-seq data sets demonstrate the utility of the package for detecting EndoMT-associated transcriptional programmes, comparing EndoMT and Hallmark EMT enrichment, together with marker subtype-level exploratory analysis of partial EndoMT states.

To our knowledge, this represents the first Hallmark-style EndoMT-focused gene set developed as a general-purpose transcriptomic enrichment resource. It addresses an important gap, as EndoMT has historically been considered within the broader framework of EMT rather than as a distinct endothelial-specific biological process (Carew, Wang and Kantharidis, 2012). With increasing recognition of pathological EndoMT in tissue fibrosis, vascular dysfunction and cancer (Qian *et al*., 2025), alongside the rapid expansion of transcriptome-scale research (Manzoni *et al*., 2018) and emergence of disease- and tissue-specific EndoMT molecular signatures (Bronson, Lyu and Xiong, 2023; Slenders *et al*., 2025; Monteiro *et al*., 2021; Lebas *et al*., 2024; Liang *et al*., 2025; Park *et al*.), there is a need for a broad, transferable gene set that captures shared EndoMT biology across vascular beds and pathological settings. Packaging this as an accessible Hallmark-style resource enables its routine application in transcriptomic enrichment analyses.

The EndoMT gene sets were structured to support both broad GSEA and more targeted exploratory interrogation of specific EndoMT-associated components. Each gene was assigned to one of seven marker subtypes, with definitions provided within the package and Supplementary Data Sheet 1 to assist interpretation of enrichment results. This structure allows users to retain or filter marker subtypes within their own workflows, tailoring analyses to their biological question, tissue context or disease model. This is useful for EndoMT, where transcriptional programmes may reflect partial or transitional states rather than a complete mesenchymal endpoint. Canonical blood and lymphatic endothelial cell markers represent genes widely accepted as lineage-defining for vascular or lymphatic endothelium, whereas endothelial-associated markers capture a more heterogeneous set of endothelial-expressed genes that vary according to vascular bed, activation and metabolic state, angiogenic and proliferative activity, developmental stage, inflammation and disease context, and thereby provide insight into context-dependent endothelial responses during EndoMT. Although the gene set was initially curated in the context of dermatological studies and a subset of endothelial-associated markers were informed by cutaneous single-cell transcriptomic data sets (Li *et al*., 2021; He *et al*., 2022), these genes should not be interpreted as skin-specific. For example, genes represented in the resource, including *ACKR1*, *EMCN*, *ICAM1*, *SELP*, *FGF2*, *LRG1* and *PLAUR*, have also been reported in non-cutaneous endothelial contexts (Schupp *et al*., 2021; Zhang, Yang and Gao, 2020; Hu *et al*., 2025; Haydinger *et al*., 2023; Barkalow, Barkalow and Mayadas, 2000; Bikfalvi *et al*., 1997; Dritsoula *et al*., 2022; Hong *et al*., 2019; Wang *et al*., 2013; Pepper *et al*., 1993; Breuss and Uhrin, 2012), while additional genes such as *ASS1*, *FADS2*, *HMGCS1*, *SCD* and *SQLE* show broader tissue expression (Uhlén *et al*., 2015). This supports the transferability of the gene set, while emphasising that enrichment results should be interpreted in relation to the tissue, disease model and experimental design under study. The inclusion of tumour endothelial markers broadens the potential utility of the resource for cancer transcriptomic studies. Given increasing evidence that EndoMT contributes to tumour microenvironment remodelling, progression and treatment resistance (Clere, Renault and Corre, 2020; Choi *et al*., 2020), inclusion of this subset enables interrogation of EndoMT-associated vascular changes in cancer data sets.

Overlap analysis between the curated EndoMT gene set and MSigDB Hallmark gene sets demonstrated the greatest overlap with Hallmark EMT. This was to be expected given that EndoMT and EMT share cellular plasticity and tissue-remodelling biology. Overlap with the Hallmark apical junction signature was also biologically plausible, reflecting shared transcriptional programmes with cell-cell adhesion, junctional stability, barrier integrity and architectural remodelling, all of which are relevant to endothelial identity loss in EndoMT (Lebas *et al*., 2024; Yoshimatsu and Watabe, 2022; Fang, Hultgren and Hughes, 2021). These findings support EndoMT as an EMT-related but distinct endothelial-specific process, reinforcing the rationale for a dedicated EndoMT transcriptomic framework rather than reliance on Hallmark EMT alone.

Benchmarking in the publicly available bulk RNA-seq data set GSE118446 demonstrated expected enrichment in an experimentally validated EndoMT-positive setting. In IL-1β/TGF-β2 co-treated HUVECs and HPAECs, the EndoMT UP gene set was positively enriched and the EndoMT DOWN gene set was negatively enriched, concordant with the original study’s RT-qPCR and protein-level evidence of EndoMT induction (Monteiro *et al*., 2021). This supports the use of the package for detecting coordinated EndoMT-associated transcriptional changes. Additionally, the single-ligand treated HUVEC analyses illustrated how transcriptome-wide enrichment can identify partial EndoMT-associated transcriptional states that may not be captured by focused endpoint marker-panel validation alone. This is particularly relevant because EndoMT is a dynamic and potentially reversible process, which can contribute to disease through persistent or transient intermediate endothelial-mesenchymal states, rather than through acquisition of a long-term mesenchymal fate (Fang, Hultgren and Hughes, 2021; Tombor *et al*., 2021). Given increasing interest in EndoMT as a potential therapeutic target across fibrosis, vascular disease and cancer (Man *et al*., 2019; Chen, Schwartz and Simons, 2020; Zeisberg *et al*., 2007b; Jia *et al*., 2021; He *et al*., 2013; Cipriani *et al*., 2015; Kanasaki *et al*., 2014; Chen *et al*., 2017; Choi *et al*., 2020; Yin and Wang, 2023), the present EndoMT gene set resource may help prioritise mechanistic pathways associated with early or partial EndoMT for downstream validation before a full phenotypic transition is established.

The in-house combined TGF-β1/2/3-treated HUVEC data set GSE342578 provided a second use case. Three-day high-dose (10 ng/ml) combined TGF-β1/2/3 exposure showed significant negative enrichment of the EndoMT UP gene set. Marker subtype-level analysis further demonstrated relative repression of EndoMT positive regulators, mesenchymal-like cell markers and tumour endothelial markers, while neither the EndoMT UP or EndoMT DOWN gene set was significantly enriched in the medium-(2.5 ng/ml) or low-dose (0.25 ng/ml) treatment arms. While these findings alone do not establish a mechanism, they demonstrate that the EndoMT gene sets can distinguish transcriptional states that are not directionally concordant with the established EndoMT-positive benchmark GSE118446. This is consistent with the broader concept that TGF-β-associated EndoMT induction is dose- and duration-dependent, supported by the accompanying single-cell RNA-seq (scRNA-seq) analysis reported by Monteiro et al. (Monteiro *et al*., 2021), in which time-dependent transcriptional separation was observed, with three- and seven-day treated HUVECs clustering separately. The biological basis for the response to short-duration ligand exposure observed here remains unclear in the absence of experimental validation, which is beyond the scope of this study, but may reflect early compensatory repression, negative feedback regulation or non-canonical TGF-β signalling activity (Xiao *et al*., 2015).

GO Biological Process enrichment of GSE118446 leading edge gene subsets demonstrated the downstream interpretive value of the EndoMT gene set resource. This post-GSEA analysis showed that while Hallmark EMT captured broad stromal, extracellular matrix and developmental processes, EndoMT UP highlighted additional migratory and vascular-remodelling biology, with the GO term “angiogenesis” showing consistently strong enrichment across all contrasts. This is biologically consistent with the overlap between transcriptional programmes regulating angiogenesis and those that govern EndoMT, along with pro-angiogenic endothelial cells being increasingly recognised as a partial EndoMT phenotype (Welch-Reardon, Wu and Hughes, 2015; Welch-Reardon *et al*., 2014). EndoMT UP also captured inflammation-associated biology in alignment with reported EndoMT literature. In chronic inflammation, persistent endothelial activation and increased leukocyte adhesion, amongst other factors, leads to endothelial dysfunction and pathological EndoMT (Yoshimatsu and Watabe, 2022). Our findings also complement previous transcriptomic analyses of EndoMT models. Namely, GO analysis of four-day IL-1β/TGF-β1 co-treated HUVECs, human arterial endothelial cells (HAECs) and human pulmonary microvascular endothelial cells (HPMVECs), suggested that EndoMT may act as a checkpoint for leukocyte chemotaxis, adhesion and transendothelial migration (Bronson, Lyu and Xiong, 2023). Taken together, these findings support the ability of the EndoMT gene sets to resolve endothelial-relevant mesenchymal programmes beyond Hallmark EMT alone.

Benchmarking in the publicly available scRNA-seq data set GSE237895 provided an opportunity to assess the curated EndoMT gene set in an alternative transcriptomic data type and across single-cell profiling-derived endothelial cell subpopulations undergoing transdifferentiation. These subpopulations are representative of the phenotypic heterogeneity observed across graded EndoMT vascular remodelling (Lebas *et al*., 2024). Most notably, chronic TGF-β1 stimulation and SNAI1 overexpression showed significant positive enrichment of the EndoMT UP gene set in the mesenchymal-like endothelial cell subpopulation, along with non-significant negative enrichment of the EndoMT DOWN signature. These findings complement the enrichment patterns observed in the former chronic TGF-β exposure model, in which GSE118446 IL-1β/TGF-β2 co-treated HUVECs and HPAECs also demonstrated significant positive enrichment of the EndoMT UP gene set and significant negative enrichment of the EndoMT DOWN gene set. Additionally, the human metastatic melanoma scRNA-seq data set GSE72056 provided a tissue-derived model of the heterogenous tumour microenvironment, in which to assess the ability of the EndoMT gene sets to distinguish distinct endothelial and mesenchymal/stromal transcriptional states, supporting their utility beyond treatment-based *in vitro* models. These findings illustrate how the curated EndoMT gene sets can be applied to transcriptional analyses of complex tumoural microenvironments and are conceptually transferable to other heterogenous EndoMT-relevant disease contexts.

Several limitations should be acknowledged when applying the package. First, the EndoMT gene sets were curated through a comprehensive narrative literature review rather than a formal systematic review. Although a formal systematic review may have provided more structured evidence weighting for individual gene inclusion or redundancy reduction, this approach reflects the intended scope of the resource as a curated, biologically informed EndoMT gene set, rather than a computationally derived signature equivalent to MSigDB Hallmarks. Second, some gene assignments to EndoMT UP or EndoMT DOWN were manually allocated based on related endothelial and mesenchymal biology in the absence of direct validation in EndoMT-specific systems. This reflects a broader challenge in EndoMT biology, where gene expression is heterogeneous, context-dependent and influenced by vascular bed, stimuli, disease state and degree of transition. Gene sets were therefore orientated around anticipated expression during partial EndoMT rather than full EndoMT, as many EndoMT-relevant genes represent transitional processes that may be upregulated during early induction but subsequently maintained, attenuated or downregulated at later stages. Enrichment results derived from this resource should therefore be interpreted in relation to the user’s biological model and supported by orthogonal validation where possible. A corresponding interpretive disclaimer is included within the R package documentation. Third, the benchmarking analyses used data sets that differed in treatment, duration, transcriptomic platform and upstream differential expression processing, so biological comparisons between data sets should be interpreted cautiously as these differences in data type, preprocessing, statistical framework and differential expression may influence ranked gene lists and downstream enrichment results. However, the aim was not to compare exact differential expression magnitudes between data sets, but to assess directional concordance or discordance of EndoMT gene set enrichment across biologically relevant models, using the same downstream GSEA workflow.

A major strength of this resource is its accessibility and interoperability. The EndoMT R package distributes the curated gene sets as standard GMT files, a plain-text format that can be accessed directly within R, but can also be exported for use in other GMT-compatible genomics, pathway and network analysis environments, for example Cytoscape/Enrichment Map, GenePattern and g:Profiler/gprofiler2 (Merico *et al*., 2010; Reich *et al*., 2006; Kolberg *et al*., 2020). In addition, the package provides exported R objects, gene-level annotation metadata and marker subtype classifications, enabling users to implement EndoMT-focused enrichment analysis without manually reconstructing the gene sets. Although benchmarking in this study was performed using bulk and single-cell RNA-seq data, the underlying principles of gene set enrichment and GMT-based gene set representation are transferable to other transcriptomic data types, including microarray and spatial transcriptomics, with appropriate preprocessing and interpretation. The inclusion of marker subtype annotations allows users to move beyond broad EndoMT UP and EndoMT DOWN enrichment results and interrogate specific components of the enrichment signal. The GMT files can also be inspected, edited and customised, providing a transparent and scalable framework that can be adapted as new EndoMT transcriptomic data sets and disease-specific signatures emerge. Future iterations could expand the resource to include mouse gene symbols and ortholog-mapped gene sets, in alignment with the cross-species structure of MSigDB (Castanza *et al*., 2023). The accompanying GitHub repository provides reproducible code and integrated GSEA result objects from the benchmarking analyses, supporting reuse, auditability and adaptation by the wider research community.

In conclusion, the EndoMT R package provides an open, extensible and interoperable resource for EndoMT-focused transcriptomic analyses. Benchmarking illustrates its ability to detect enrichment patterns associated with established, partial and divergent EndoMT-associated states, while post-GSEA GO Biological Process analysis highlights endothelial-relevant mesenchymal biology beyond MSigDB Hallmark EMT.

## 5 Conflict of Interest

The authors declare that the research was conducted in the absence of any commercial or financial relationships that could be construed as a potential conflict of interest.

## 6 Author Contributions

KB: Conceptualization; Data curation; Formal analysis; Funding acquisition; Investigation; Methodology; Software; Validation; Visualization; Writing – original draft.

JC: Conceptualization; Software; Writing – review and editing.

PEL: Funding acquisition; Project administration; Supervision; Writing – review and editing.

LW: Conceptualization; Data curation; Formal analysis; Funding acquisition; Investigation; Methodology; Supervision; Validation; Visualization; Writing – review and editing.

JA: Conceptualization; Project administration; Supervision; Writing – review and editing.

## 7 Funding

This research was supported by Cancer Research UK through a pre-doctoral bursary (RCCPDB-Nov22/100004; to KB) and a Clinical Academic Training Fellowship (SEBCATP-2024/100001; to KB and PEL), the North Eastern Skin Research Fund (Aug22; to KB and PEL), the John George William Patterson Foundation (Jun26/NU-026800; to LW, PEL, JA), and the NIHR Biomedical Research Centre (BRC): Newcastle awarded to The Newcastle upon Tyne Hospitals NHS Foundation Trust, Faculty of Medical Sciences, Newcastle University and Cumbria, Northumberland, Tyne and Wear NHS Foundation Trust (to KB, LW and PEL).

## 8 Data Availability Statement

Publicly available data sets were analysed in this study. These data can be found at the NCBI Gene Expression Omnibus (GEO) under accessions GSE118446 [https://www.ncbi.nlm.nih.gov/geo/query/acc.cgi?acc=GSE118446], GSE237895 [https://www.ncbi.nlm.nih.gov/geo/query/acc.cgi?acc=GSE237895] and GSE72056 [https://www.ncbi.nlm.nih.gov/geo/query/acc.cgi?acc=GSE72056]. The original contributions presented in this study will be released upon publication and can be found here: [https://www.ncbi.nlm.nih.gov/geo/query/acc.cgi?acc=GSE342578].

The EndoMT R package, a complete worked example script using the GSE118446 data set, together with processed R objects required to reproduce the integrated MSigDB Hallmark and EndoMT enrichment visualisations, are hosted in a GitHub repository [https://github.com/kbdermdoc/EndoMT]. The repository will remain private during peer review and will be made publicly available upon publication.

The curated EndoMT gene set metadata, GMT file contents, MSigDB Hallmark gene sets used for overlap analysis, DESeq2 results, TERM2GENE files, leading edge gene lists and GO Biological Process enrichment results are included in the Supplementary Material. All supplementary data sheets and supplementary table 1 will remain private during peer review and will be made publicly available upon publication.

---

**Supplementary Figure 1.**
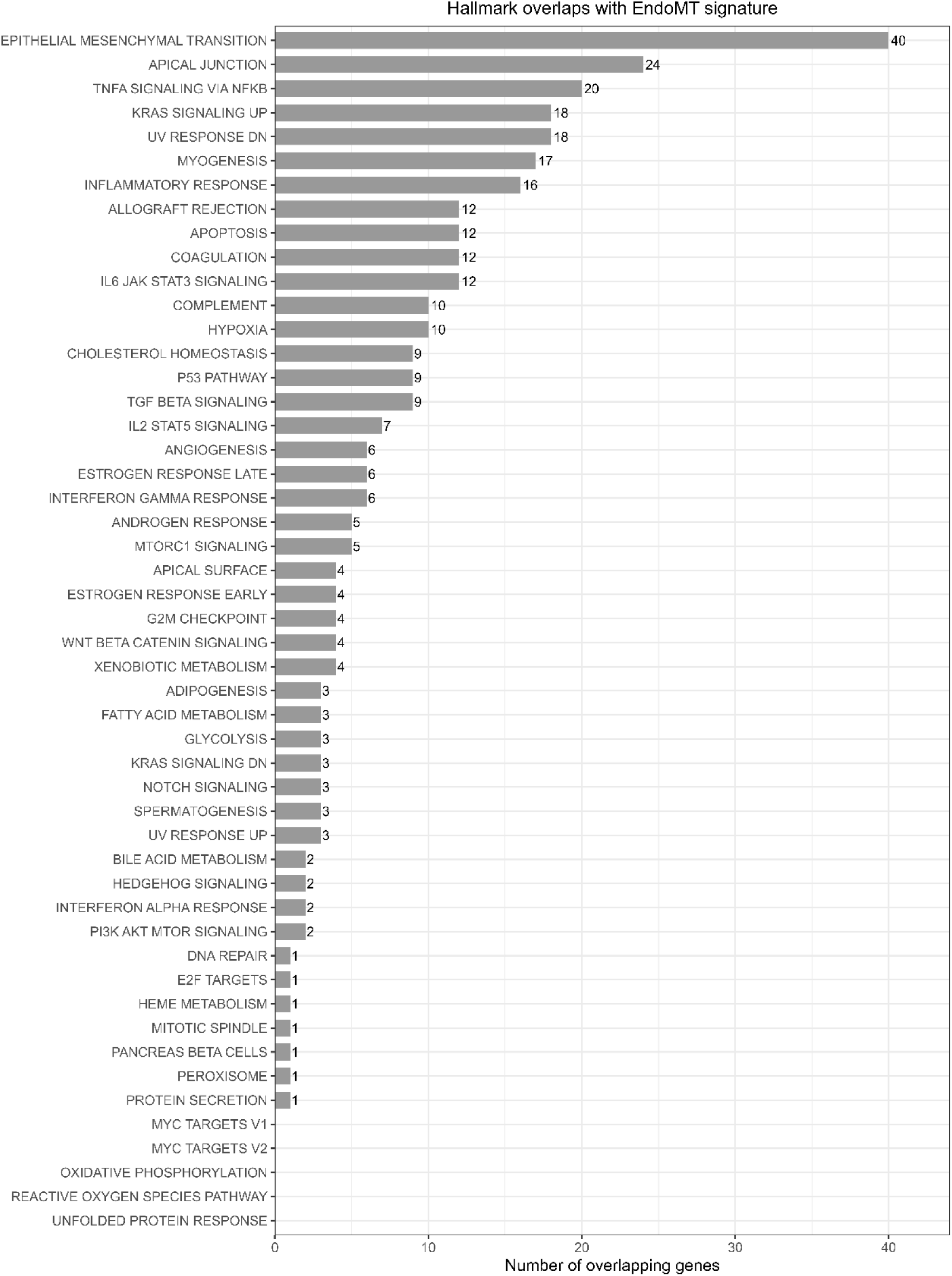
Absolute gene overlap between EndoMT and MSigDB Hallmark gene sets. Bar plot showing the number of genes shared between the curated EndoMT gene set and each MSigDB Hallmark gene set.

**Supplementary Figure 2.**
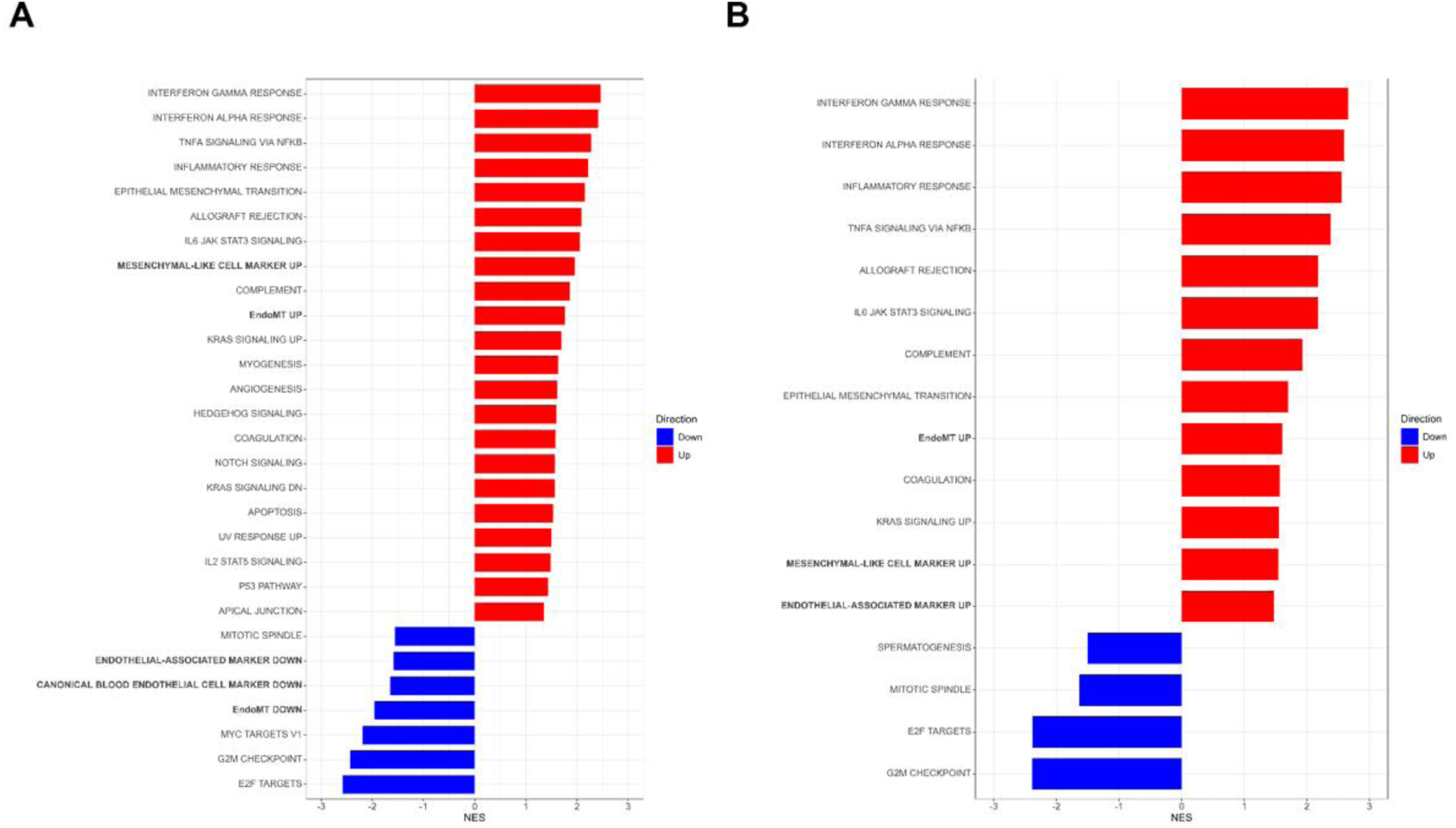
GSEA of curated EndoMT marker subtypes supports EndoMT induction in GSE118446 HUVECs and HPAECs following IL-1β/TGF-β2 co-treatment. GSEA was performed at the level of curated EndoMT marker subtypes in GSE118446 (A) HUVECs and (B) HPAECs treated with combined IL-1β/TGF-β2 for seven days compared with untreated controls. GSEA result objects from EndoMT R package-derived marker subtype-specific EndoMT UP and EndoMT DOWN gene sets, together with MSigDB Hallmark gene sets, were integrated prior to visualisation. NES bar plots show significantly enriched gene sets in HUVECs and HPAECs. Only gene sets passing BH-adjusted P value ≤ 0.05 are shown.

**Supplementary Figure 3.**
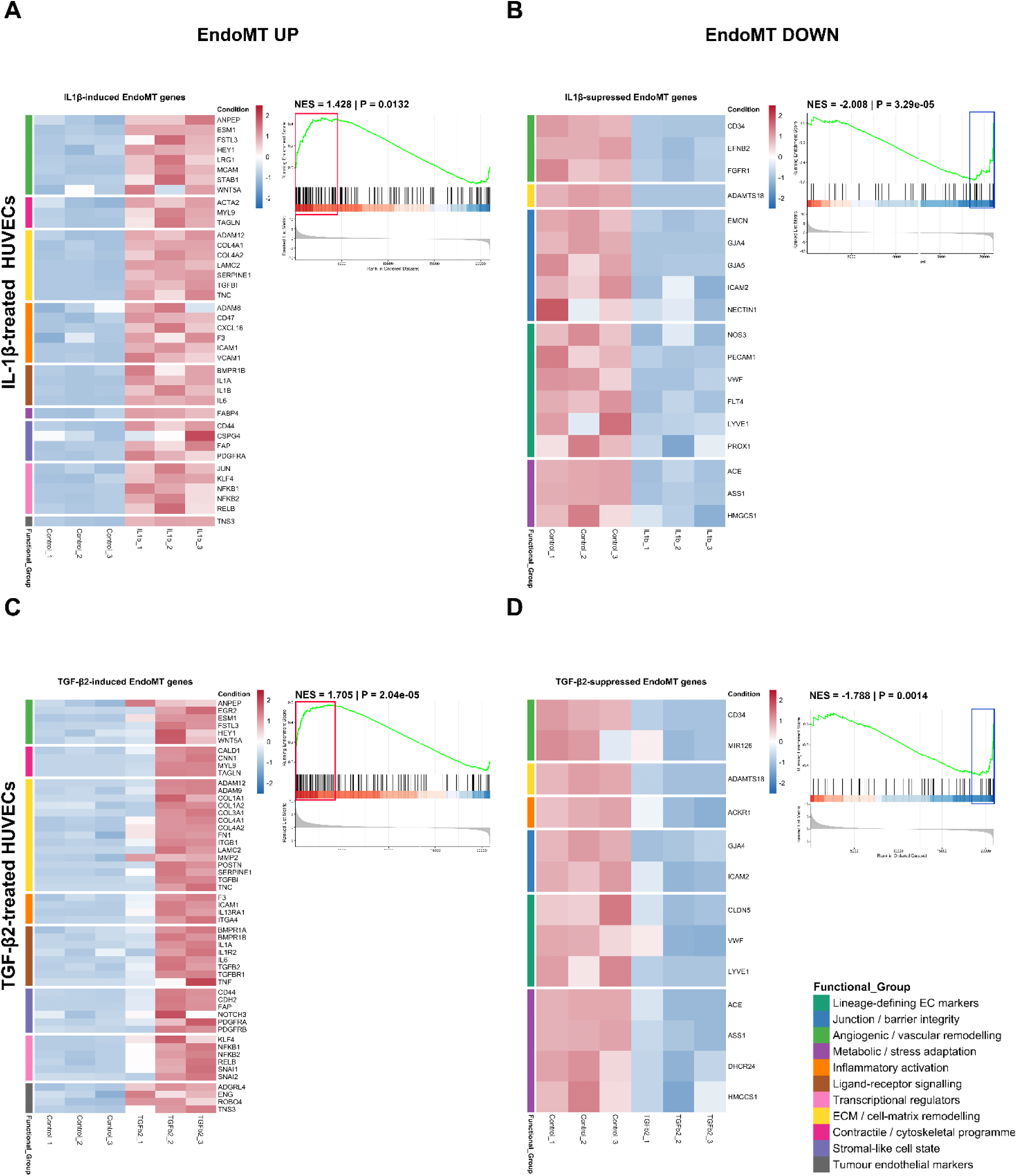
Leading edge analysis of EndoMT UP and EndoMT DOWN gene set enrichment in GSE118446 single ligand-treated HUVECs. Enrichment plots and corresponding heatmaps of the leading edge genes grouped by functional annotation for (A, B) IL-1β-treated and (C, D) TGF-β2-treated HUVECs.

**Supplementary Figure 4.**
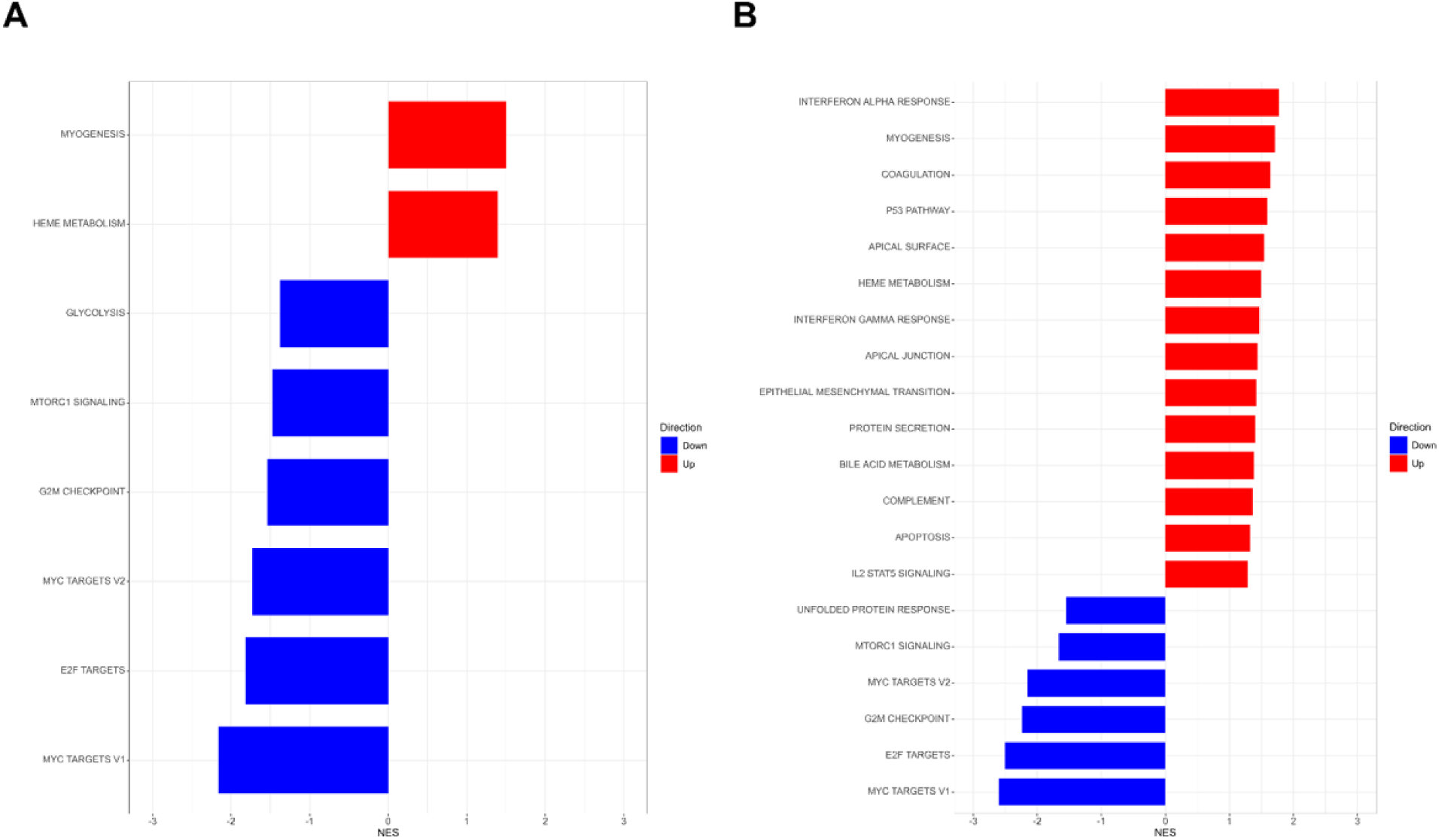
Low- and medium-dose combined TGF-β1/2/3 exposure does not induce EndoMT gene set enrichment in HUVECs. GSEA was performed on in-house GSE342578 HUVECs treated with 0.25 ng/ml (low-dose) and 2.5 ng/ml (medium-dose) combined TGF-β1/2/3 for three days compared with untreated controls, using EndoMT R package-derived gene sets, alongside MSigDB Hallmarks. NES bar plots of EndoMT UP and EndoMT DOWN gene set enrichment, showing no significant enrichment in (A) 0.25 ng/ml (low-dose) and (B) 2.5 ng/ml (medium-dose) combined TGF-β1/2/3-treated HUVECs. Only gene sets passing BH-adjusted P value ≤ 0.05 are shown.

**Supplementary Figure 5.**
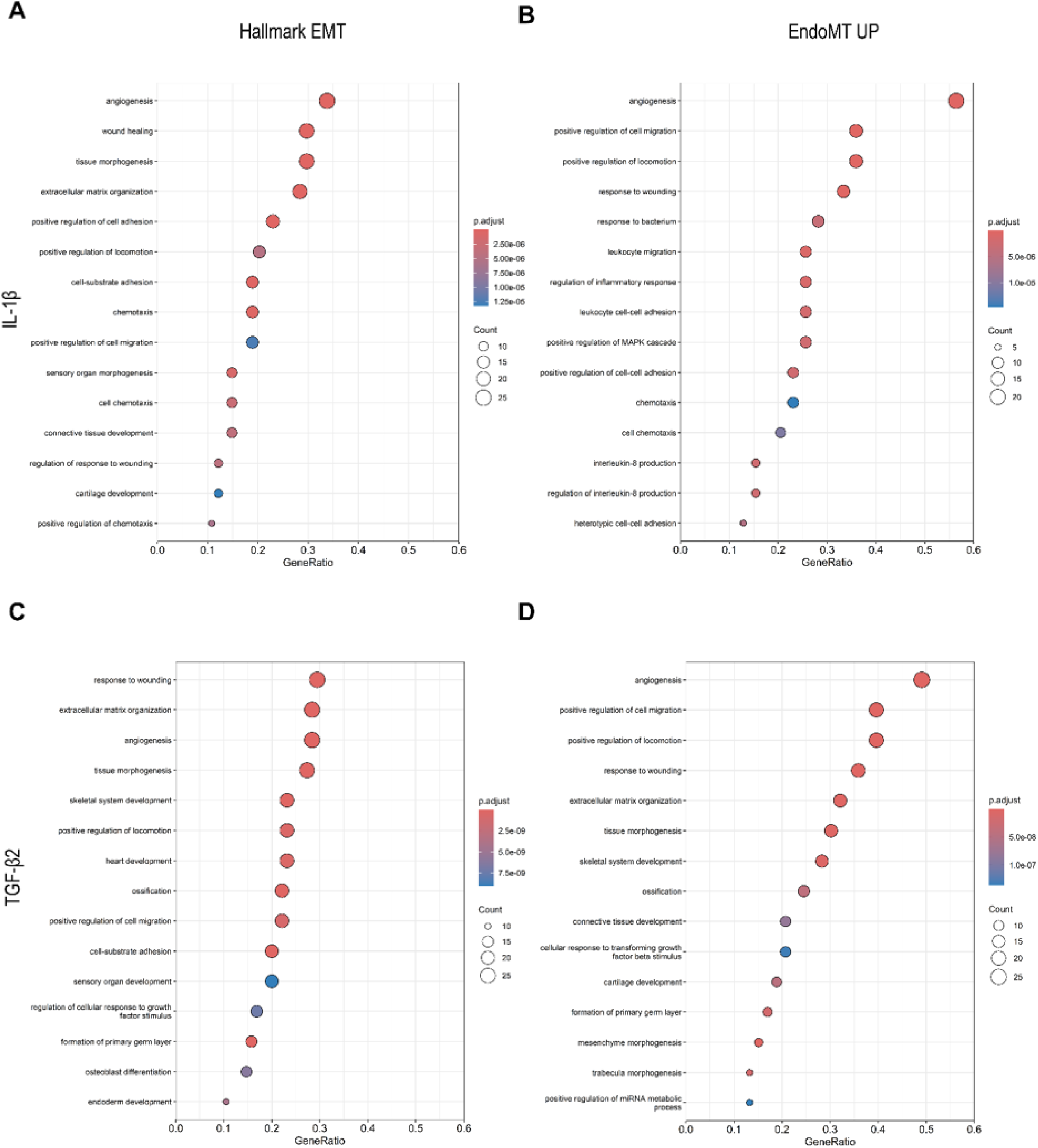
GO Biological Process enrichment of Hallmark EMT and EndoMT R package-derived EndoMT UP leading edge genes in GSE118446 single ligand-treated HUVECs. GO Biological Process enrichment analysis was performed on leading edge gene subsets derived from significantly positively enriched Hallmark EMT and EndoMT UP gene sets in GSE118446 HUVECs treated with IL-1β or TGF-β2 for seven days compared with untreated controls. Enrichment plots are shown for Hallmark EMT and EndoMT UP leading edge genes in (A, B) IL-1β-treated and (C, D) TGF-β2-treated HUVECs. GO terms were filtered by BH-adjusted P value ≤ 0.05.

**Supplementary Figure 6.**
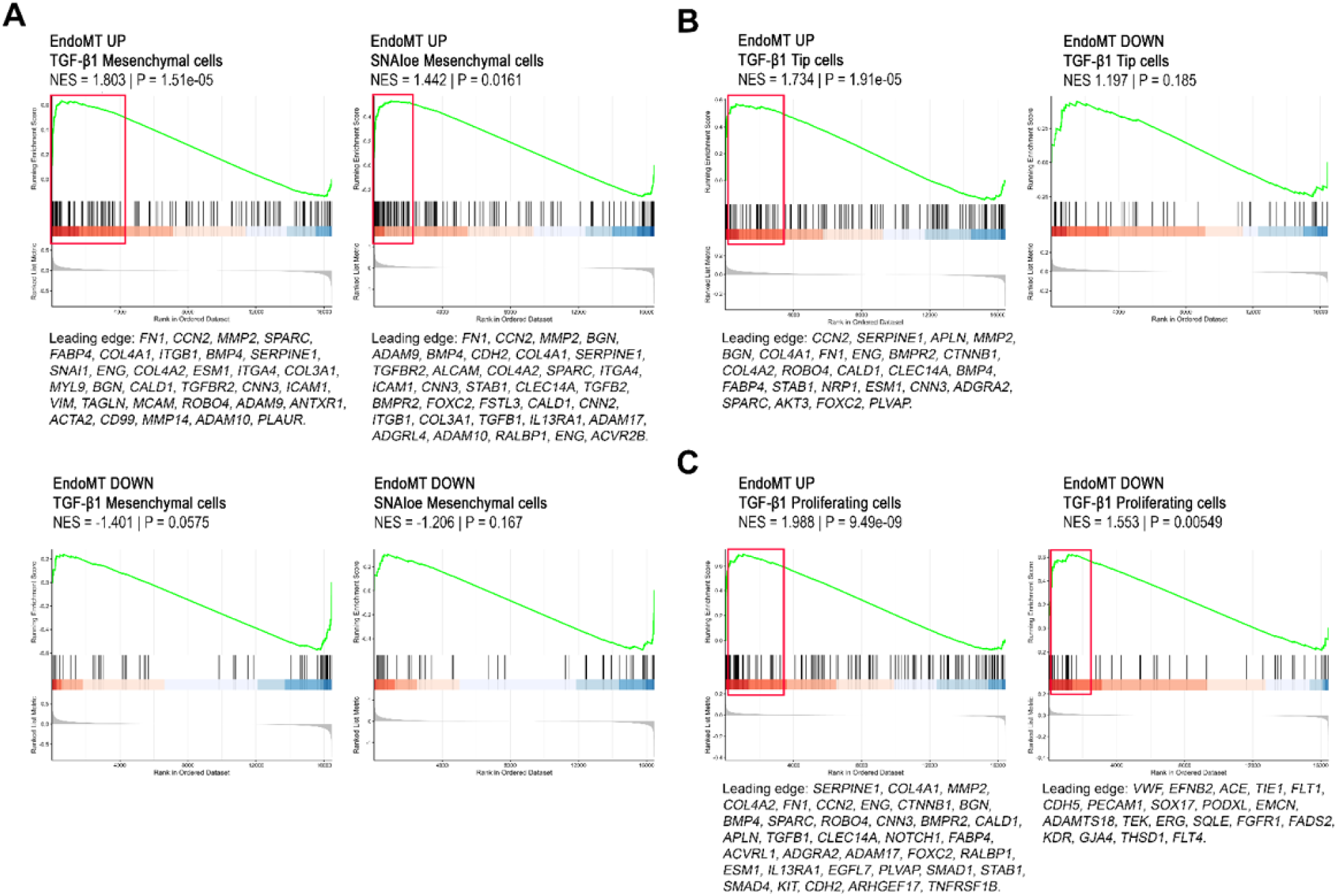
Application of EndoMT R package gene sets capture TGF-β1 and SNAI1-associated EndoMT programmes in GSE237895 single-cell endothelial subpopulations. GSEA was performed on GSE237895 endothelial cell subpopulations derived from seven-day TGF-β1-treated or SNAI1 overexpressing (SNAI1oe) HUVECs, compared with untreated controls, using EndoMT R package-derived EndoMT UP and EndoMT DOWN gene sets, alongside MSigDB Hallmark gene sets. EndoMT UP enrichment plots show significant positive enrichment in (A) TGF-β1-treated and SNAI1oe mesenchymal-like endothelial cells, (B) TGF-β1-treated tip cells and (C) TGF-β1-treated proliferating cells. Corresponding leading edge genes are displayed for all significant gene set enrichment results (BH-adjusted P ≤ 0.05).

**Supplementary Figure 7.**
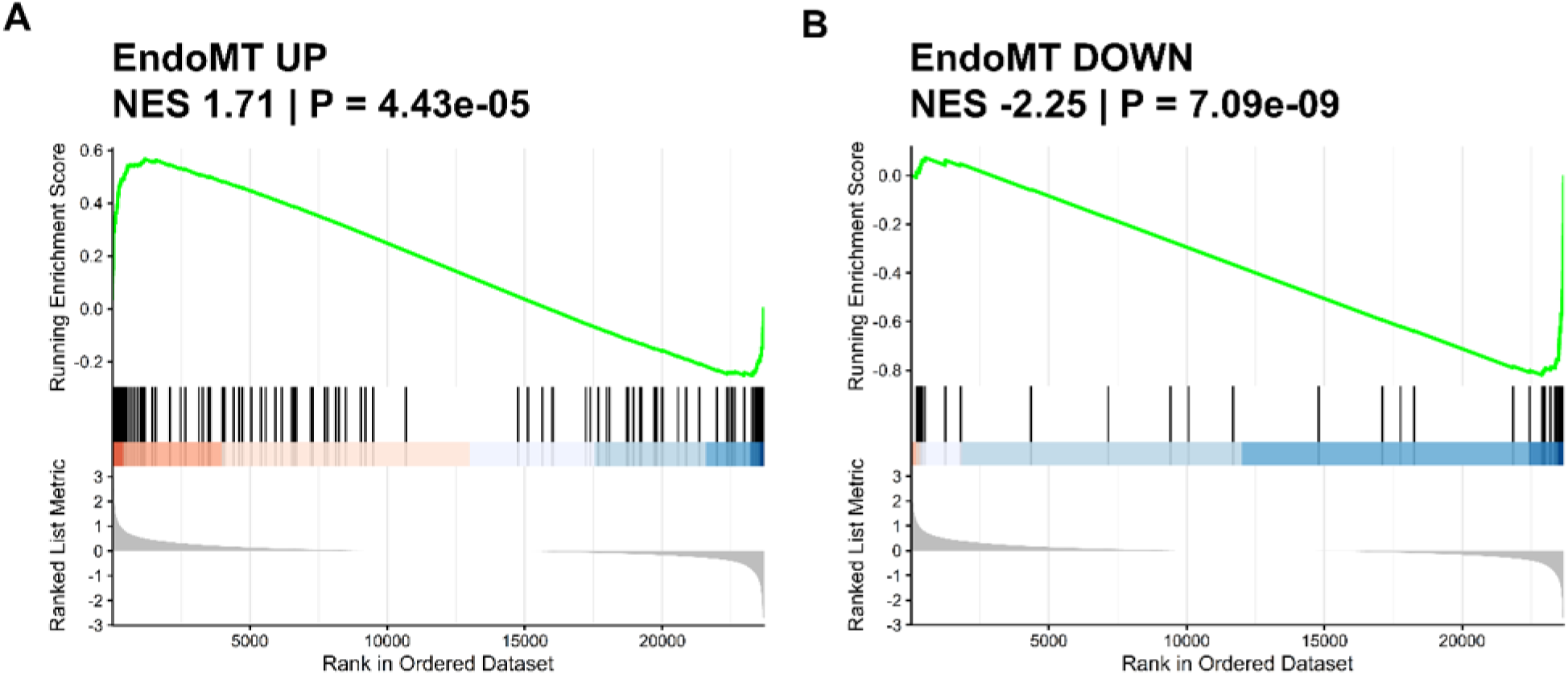
EndoMT gene sets discriminate mesenchymal and endothelial states in GSE72056 human metastatic melanoma single-cell subpopulations. GSEA was performed comparing CAF and endothelial cell subpopulations in GSE72056 using EndoMT R package-derived (A) EndoMT UP and (B) EndoMT DOWN gene sets.

## Notes

### Competing Interest Statement

The authors have declared no competing interest.

